# The Functional and Genetic Architecture of Olfaction in Deer Mice

**DOI:** 10.1101/2025.05.12.653383

**Authors:** Jean-Marc Lassance, Andreas F. Kautt, Landen Gozashti, Hopi E. Hoekstra

## Abstract

Mammals have well-developed olfactory systems, but the molecular logic driving the extraction of biological information, how this logic evolves and contributes to differences in behavior remain unclear. To address this, we generated a chromosome-level assembly of the North American deer mouse, *Peromyscus maniculatus*, meticulously annotating the complex olfactory subgenome. Comparative analysis revealed high evolutionary plasticity within chemosensory gene families. Rodent olfactory and vomeronasal sensory pathways, predominantly associated with odor learning and innate responses, respectively, show divergent evolutionary trajectories–one conserved, the other rapidly evolving. Despite overall vomeronasal gene dynamism, we identified receptors conserved across evolutionary timescales. Using the activation of these receptors as readouts, we demonstrate the conservation of receptor-ligand pairings for subsets of vomeronasal sensory neurons. Collectively, our findings provide insights into the evolution of the mammalian olfactory systems, highlighting key differences in receptor repertoires and establishing deer mice as a model clade to investigate neural mechanisms underlying the evolution of olfactory pathways and their impact on behavioral evolution.

**Teaser:** The deer mouse genome analysis reveals conserved and divergent components of the rodent olfactory subgenome.

## Introduction

Animals possess exquisitely sophisticated sensory systems which allow them to construct internal representations of the world, determine appropriate actions, and navigate the complexity of diverse environments and social contexts. Olfaction –the sense of smell– orchestrates the detection of odors in which animals are immersed within their natural environments. These odors –and the information they convey– permit to locate food sources, identify potential mates, avoid predators, etc. among an array of ecologically essential functions. Thus, olfaction plays a major role in organismal fitness, both for survival and reproduction (*1*). Essentially, the olfactory system acts as an odor-analyzer that extracts biological information and ascribes a meaning to odors in the chemosensory brain, establishing the basis for behavioral responses. In mammals, this crucial function is mediated by the activation of olfactory sensory neurons housed in anatomically and molecularly distinct sensory epithelia, namely the main olfactory epithelium (MOE) and the vomeronasal organ (VNO) (*2*). Olfactory sensory neurons each sample a portion of chemical space, defined by the type of stimuli they perceive, the molecular properties of these stimuli, and the neural circuits with which they are associated beyond their axonal projections in the olfactory bulb (*3*). Odors have traditionally been divided into two broad categories. Most odors convey information about the wider world, and the behavioral responses these elicit can be shaped by experience and learning, allowing animals to adjust their behavior to the ever-changing environment surrounding them. In parallel, a distinct class of chemical cues, pheromones and kairomones, represent both temporal and spatial information that animals can evaluate before executing the appropriate innate, genetically pre-programmed behavioral sequences (*1*). The olfactory and vomeronasal systems, and the distinct neural pathways to which they convey information, are associated with this dichotomy (*4*). Yet, despite segregated connectivity, both systems overlap partly in their functions and interact in the control of mate recognition, sexual and social behaviors (*5*).

In mammals, chemoreception is orchestrated by a diverse array of multi-gene families mainly encoding G protein-coupled receptors (GPCRs)(*6–11*): odorant receptors (Olfr)(*6*), vomeronasal receptors type 1 and 2 (Vmn1r and Vmn2r)(*7, 8*), trace amine-associated receptors (TAAR)(*9*), formyl peptide receptors (FPR)(*10*), and membrane-spanning 4-domains subfamily A (MS4A)(*11*). This diversity highlights the challenge posed by the complexity of odorants, which can only be tackled by possessing a large repertoire of diverse olfactory receptors. Comparative genomic studies revealed that these multi-gene families vary in size due to the stochasticity of birth-death evolutionary processes as well as adaptive evolution(*12*), yet we know little about the functional consequences of the many changes in chemosensory receptor repertoires revealed by comparative genomic studies. Part of this is due to a limited understanding of the molecular logic by which the olfactory system encodes the information contained in the environment, and how this logic changes during evolution and adaptation to constitute the substrate for behavioral evolution. With new insights into the basic logic of mammalian olfaction continuing to be discovered (*11*), we are beginning to have a solid understanding of the olfactory systems in *Mus musculus*. However, we know less whether these patterns are the same in other rodent species or how they evolve in response to differences in ecological community and social environment. This advocates for the examination of additional model species.

Deer mice (genus *Peromyscus*) are approximately 25 millions of years diverged from the house mouse (*13*) and represent an exciting model clade in which to study the evolutionary origins and implications of variation in olfactory gene repertoires. The genus comprises 55 species than span remarkable ecological diversity. The most abundant of these, *P. maniculatus*, is found in diverse habitats across most of North America (*14*)(Fig 1). *Peromyscus* may be considered an example of an adaptive radiation in terrestrial mammals and constitutes one of the best studied rodent genus in terms of behavior and ecology (*15*). Because of its widespread distribution, it occurs with distinct congeners as well as predators. Moreover, within this genus, distinct mating systems have evolved: for example, while *P. maniculatus* is promiscuous and litters are most often sired by multiple males (*16*), its sister species, *P. polionotus*, is both socially and genetically monogamous (*17*). Thus, *Peromyscus,* in general, and *P. maniculatus,* in particular, provides an alternative to common rodent models for studying the neural mechanisms underpinning social and non-social olfactory recognition and the associated behaviors (*18*).

**Fig. 1.**
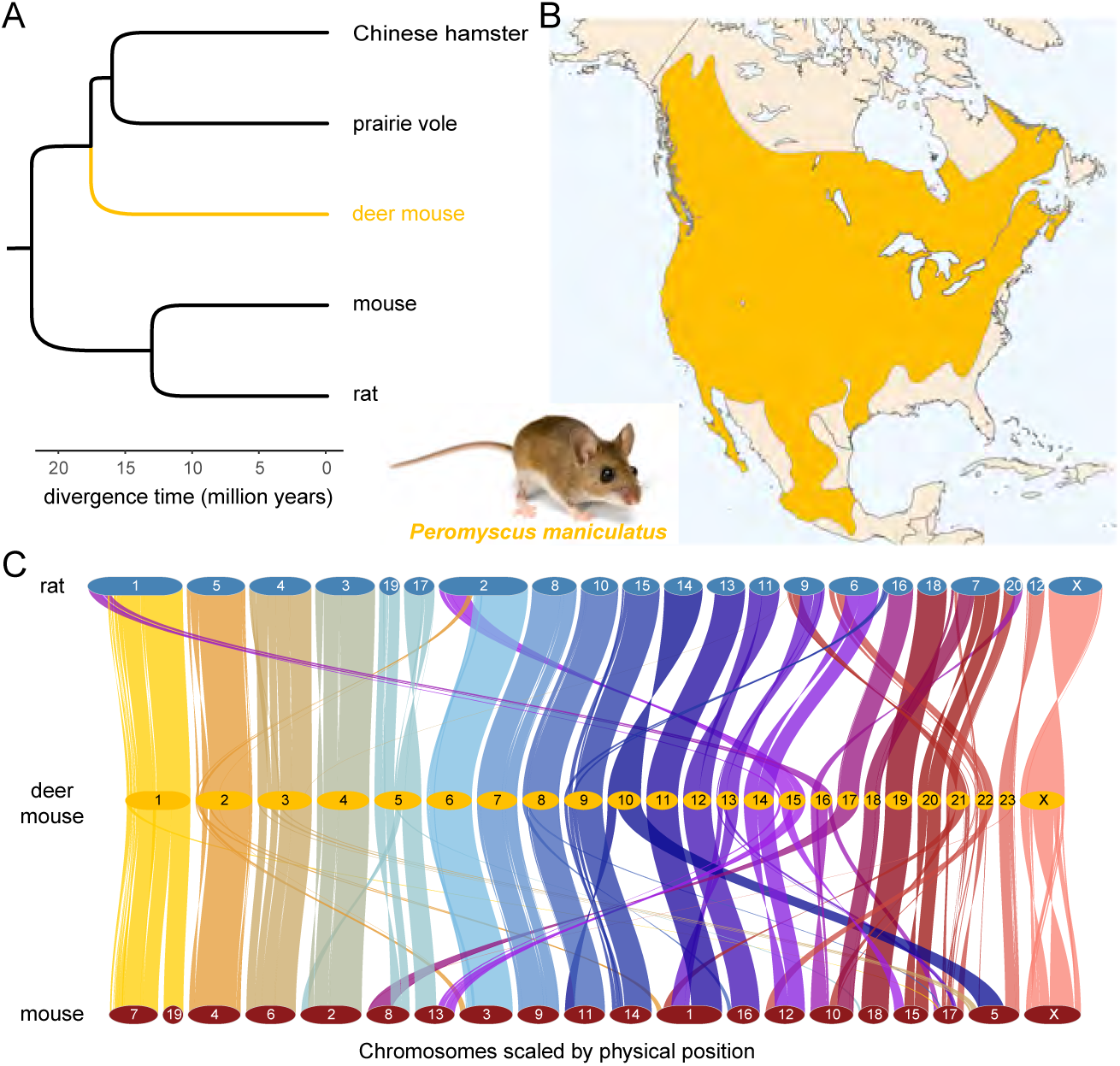
Nuclear genome assembly for the prairie deer mouse. (**A**) Simplified phylogeny showing the relationships among muroid rodent model organisms. (**B**) Distribution of *P. maniculatus*, which is found throughout most of North America and inhabits most biomes on the continent. (**C**) Syntenic network of orthologous regions between deer mouse, rat, and mouse genomes. Ribbons are color-coded by synteny to deer mouse chromosomes.

Here we report on a chromosome-level assembly of the *P. maniculatus* genome, which has allowed the further integration of ecological, evolutionary, behavioral, and genomic approaches with this non-traditional classic model species. Our comparative approach corroborates that olfactory receptor genes are among the most volatile genes in the genome. However, our characterization of the olfactory subgenome of *Peromyscus* reveals radically different trajectories followed by olfactory receptor gene families, from substantial conservation in the repertoire of genes expressed by neurons of the main olfactory system to limited overlap in vomeronasal system repertoires across species. Finally, our study uncovers a set of receptors involved in social communication and semiochemical detection, providing a molecular foundation to further dissect the evolution of neural circuits related to these processes in rodents.

## Results

### Chromosome-level genome assembly by integration of Illumina assembly and genetic maps

We assembled the deer mouse genome using a combination of Illumina short-read data sequenced from libraries of complementary insert sizes (Table S1). We obtained a 2.517 Gb assembly with an N50 contig size and an N50 scaffold size of 30.1 kb and 13.86 Mb, respectively (Table S2). In addition, we capitalized on the availability of four high-density genetic linkage maps from previous studies aiming at characterizing the genetic architecture of various morphological and behavioral traits in *Peromyscus* (*19–22*). Using these maps, we could assign and position 97% of the *de novo* assembled bases into chromosomes (Table S3). The total number of chromosomes covered is 24, including the X chromosome, consistent with cytogenetic studies (*23*). We assigned chromosome names based on previous reports from interspecific reciprocal whole chromosome painting, which have allowed to assign linkage groups containing known genes to *Peromyscus* chromosomes (*24, 25*).

We next assessed the completeness of the genome assembly using BUSCO (Benchmarking Universal Single-Copy Orthologs). Our estimates indicate that this assembly contains 97.6% and 96.3% of core mammalian and euarchontoglire genes, respectively (Fig. S1). The observed level of completeness and the assignment of chromosomal locations to most of the assembled bases are indicative of a high-quality reference genome, making it suitable for a range of applications, including evolutionary analyses and comparative studies (Fig. S2). Pairwise alignment of the deer mouse genome to the reference genomes of mouse and rat revealed large-scale synteny, as well as several inter-chromosomal rearrangements, consistent with observations in other mammals (*26*)(Fig. 1, Fig. S3, Fig. S4). These analyses show that the macro-organization of the *Peromyscus* genome appears more similar to *Rattus* than *Mus* (Fig. S5). This corroborates earlier reports based on low-density linkage maps (*24, 27*). The comparison of these three genomes provides an overview of the genomic regions that are conserved and play a role in genome stability as well as the regions prone to breakage and structural rearrangements.

### Comparative annotation reveals similarities and differences with house mouse

To annotate genes, we used an annotation strategy making use of multiple genome alignments and an existing high-quality annotation set (*28*). While permitting the discovery of novel genes via *ab initio* gene modeling, this approach allows to identify orthology relationships readily and with high accuracy. Encouraged by the high level of synteny revealed by our analysis, we annotated the deer mouse genome using the genome of the house mouse *Mus musculus* (GRCm38/mm10) and the high-quality and well-curated GENCODE VM15 as the reference gene/transcript set, as well as extensive transcriptome sequences from six tissues (brain, testis, skin, hypothalamus, main olfactory epithelium, and vomeronasal organ). We generated an alignment including also the rat genome (rn6) and the prairie vole genome *Microtus ochrogaster* (MicOch1.0), the latter representing an evolutionary intermediate between the mouse/rat and deer mouse lineages (Fig. 1). Because the annotation is obtained in a comparative context, we could unambiguously identify orthologs for 18,529 (84.2%) and 562 paralogs of 22,014 protein-coding genes in the *Mus* genome. In addition, *ab initio* gene prediction resulted in the detection of 10,438 putative protein-coding genes, of which 1,882 have homologies with known proteins from curated databases, suggesting that the remainder may well result from false positives among the *ab initio* predictions (*29*). Next, we evaluated the completeness of our annotation. Our BUSCO analyses indicate that our deer mouse annotation contains 98.4 % and 98% of core mammalian and euarchontoglire protein-coding genes, respectively, which is on par with the statistics obtained with the genome mode (Fig. S6). Among non-coding genes, we found a variable degree of homology, from poor in the case of immunoglobulin V genes (∼ 8%) and processed and unprocessed pseudogenes (27.9% and 20.8%, respectively) to high with lincRNAs (79.9%) and antisense RNAs (95.1%) (Table S4).

Taken together, these results indicate that protein-coding genes are well annotated in the deer mouse assembly, paving the way to identify which genes contribute to critical aspects of deer mouse biology.

### Volatility of subgenome responsible for sensory perception, pheromone reception and immunity

The *Mus* and *Peromyscus* lineages shared their last common ancestor approximately 25 million years ago (*13*). Despite superficial similarities, they exhibit marked differences in physiology and behavior which are rooted in their genomes. To capture some of the genetic features underlying the adaptive evolution of deer mice, we performed two sets of analyses. First, we examined the functions of protein-coding genes for which homologs could not be detected in the deer mouse genome during the *Mus* gene annotation transfer. We note that genes missing from the comparative annotation transfer could result from different scenarios. A first group may comprise genes which evolved via lineage-specific gene duplication and retrotransposition events after the split of their respective lineages. In this case, the duplicated regions do not exist in the other lineage. Another possibility is that the regions corresponding to those genes were lost as a consequence of structural changes caused by the numerous intra- and inter-chromosomal rearrangements revealed by our analysis of synteny and what is known of the evolution of chromosomes in rodents (*30*). Finally, these may correspond to rapidly evolving genes and genomic regions that diverged to such an extent that homology can no longer be detected due to failure of similarity-search approaches to correctly identify such regions (*31*). We note, however, that our whole-genome alignment-based annotation transfer strategy should be more robust, yet not immune, to this phenomenon. Our analysis indicates that the ‘depleted’ genes for which homologs could not be detected are mainly associated with regions with no correspondence in the *P. maniculatus* genome (Fig. S7). This suggests that structural variation, and the associated loss of gene copies or sequences, has been a major contributor in the differences in gene contents between these lineages. To investigate the type of functions fulfilled by these genes, we performed a gene ontology enrichment analysis with the *Mus* genes appearing missing (*sensu lato*) from the deer mouse annotation. The results revealed an enrichment for chemosensory perception (i.e., sensory perception of smell, response to pheromone), genes related to defense/immunity, and infection response (Fig. S8). These findings recapitulate earlier work suggesting that olfactory and immune-related genes are among the fastest-evolving genes in mammals, likely due to their roles in adaptation to changing environmental cues and pathogen coevolution (*32, 33*).

Second, we performed gene family expansion and contraction analyses using CAFE (Computational Analysis of gene Family Evolution; (*34*)). For these, we used the full complement of deer mouse protein-coding genes from our *ab-initio* and comparative annotation approach. CAFE uses a stochastic birth-death model of gene family evolution to infer the size of ancestral families. These analyses provide a genome-wide overview of families that have increased or decreased among mammalian taxa (Fig. S9A). From a total of 17,370 tested gene families, we detected 169 fast-evolving gene families/orthogroups with significant gene gain or loss events in the deer mouse (*P<* 0.05). Among 42 significantly expanded gene families were genes encoding olfactory and vomeronasal receptors (annotated for a role in G protein-coupled receptor signaling pathway, response to pheromone and sensory perception of chemical stimulus), as well as proteins involved in adaptive and innate immune responses (Fig. S9B; Supplementary Table 1). The predominant functions of the 127 families that underwent contraction were translation, signal transduction, sensory perception of smell as well as defense response (Fig. S9B; Supplementary Table 2). Remarkably, we also noted the reduction or absence of families encoding or carrying pheromone signals in *Mus*, namely major urinary proteins (MUPs), secretoglobins (SCGBs), and exocrine-gland secreting peptides (ESPs) (Fig. S10).

Altogether, these results pointed to the types of genes and biological functions that may be more susceptible to change during the evolution of deer mice, highlighting genes associated with chemosensation represent some of the most rapidly evolving genes in the deer mouse lineage.

### Contrasting evolution of the chemosensory sub-genomes

The extensive variation in chemosensory genes revealed by our analyses poses the question of which specific component(s) of the olfactory repertoire are driving this signal. To investigate the evolution of olfactory sensory perception, we surveyed the gene families associated with the perception of smell, identified their homologues in the deer mouse genome, and compared these gene repertoires with those present in *M. musculus* and *R. norvegicus*. We first produced a curated annotation of olfactory receptors in *Peromyscus* using protein homologs and RNA sequencing datasets (RNA-Seq) from the whole olfactory mucosa and the vomeronasal organ, two major olfactory organs in mammals. We identified 1018 Olfrs, 15 MS4As, 10 TAARs, 179 Vmn1rs, 106 Vmn2rs and 3 Fprs (Fig. 2A). As a result of their evolution through segmental duplication (*12, 35*), chemosensory genes are organized in clusters along the genome, with an enrichment on chromosome 1, which houses approximately a third of all chemoreceptor genes (Fig. S11).

**Fig. 2.**
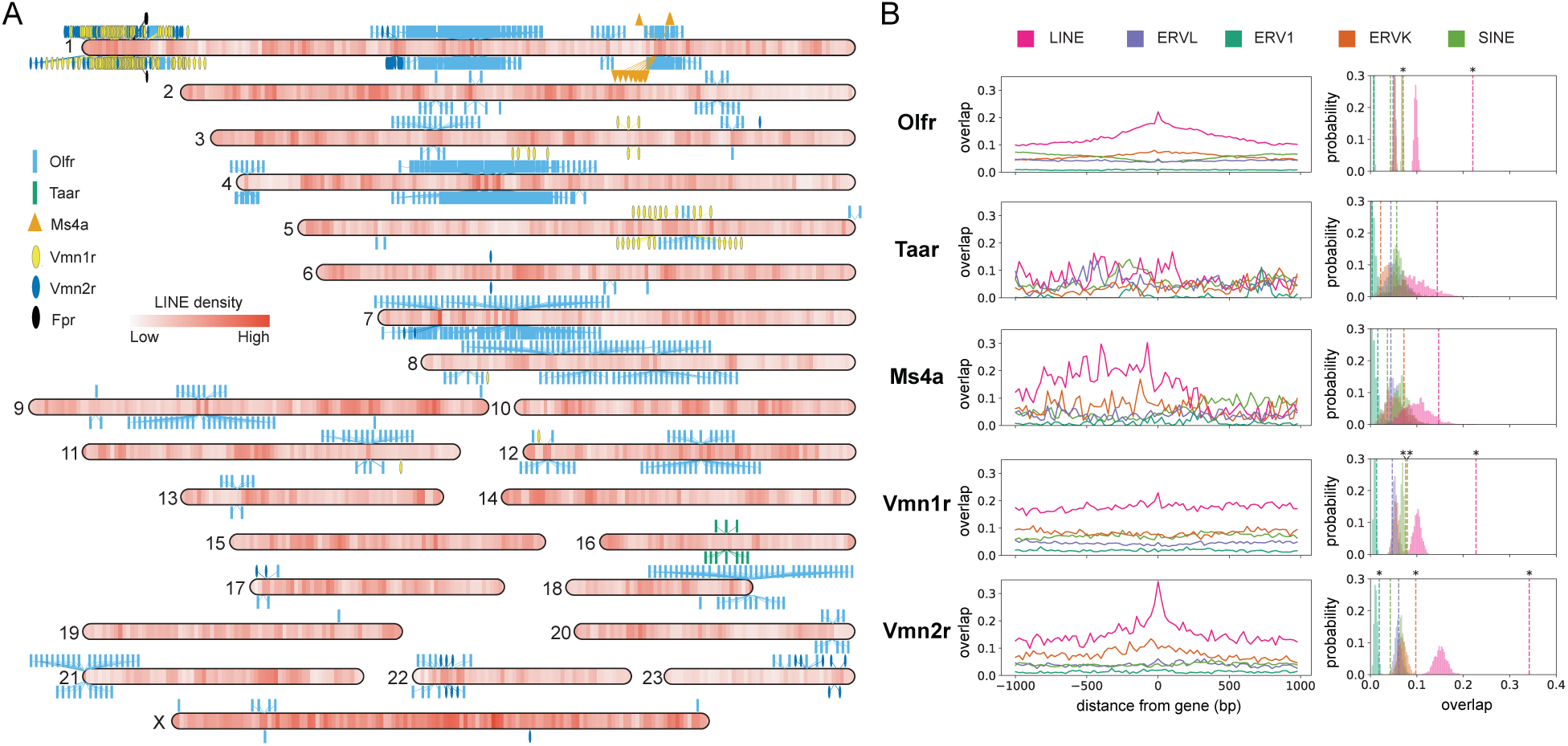
Spatial organization of the olfactory subgenome. (**A**) Chromosomal distribution of the different classes of olfactory receptor genes along the deer mouse genome. The symbols represent genes of a particular chemosensory receptor gene family, with the position indicating the gene orientation (above: forward; below: reverse). The color on a chromosome represents the LINE density. (**B**) Occupancy of individual repeat subclasses in the region flanking genes of different chemosensory receptor families. For each repeat subclass, we tested for enrichment in regions harboring olfactory receptors (dashed lines) relative to randomly resampled regions (barplots colored by repeat subclass). An asterisk marks a significant difference (1000 permutations).

Retrotransposons serve as a fertile substrate for ectopic recombination and can facilitate adaptive gene family expansion and diversification (*36–39*). At the same time, TE occupancy is not homogeneous along the chromosomes (Fig. 2A). We thus examined the association between transposable elements and the different chemoreceptor gene families (Fig. 2B), specifically the role of TEs in chemosensory gene family evolution. One prediction for TEs contributing to gene family expansion is an enrichment of TEs in gene flanking regions. To assess this possibility, we compared repeat occupancy at regions flanking chemosensory genes to the rest of the genome and performed permutation tests for enrichment of TEs relative to random expectations. We did this for LINE and SINE retrotransposons as well as the three major families of endogenous retroviral LTR elements (ERVK, ERV1 and ERVL). We observe a more than two-fold enrichment for LINEs as well as a more modest enrichment for ERVKs in the flanking regions of Olfrs, Vmn1r, and Vmn2r gene families (Fig. 2B). Interestingly, Olfrs display high levels of LINE occupancy (relative to the genome average) extending several kilobases from their breakpoints, suggesting that they also inhabit ancestrally LINE-rich regions. In contrast, Vmn1rs and Vmn2rs display steeper peaks of increased LINE occupancy, suggesting that the association between these gene families and LINEs arose more recently. Together, these results suggest that retrotransposons play a significant role in chemosensory gene family evolution in deer mice.

To place each chemosensory gene families in a broader phylogenetic context, we performed phylogenetic reconstructions using receptors from the high-quality genomes of *Mus* and *Rattus*. As gene duplications and losses occur frequently in the evolution of multigene families, one-to-one orthology might be rare or at least difficult to detect. In addition, there are limitations inherent to performing analyses using one-to-one orthologs only. To circumvent these shortcomings, we clustered genes into groups of homologous genes based on their similarity and used *orthogroup* as the unit of comparison (i.e., groups of genes that are hypothetically descended from a single ancestral gene and thus exhibit identical to highly similar function). This approach is supported by the observation that highly similar receptors perceive structurally similar molecules. Our analyses indicate that the gene complements present in *Mus* and *Rattus* often showed contrasting patterns from those seen in *Peromyscus*, oscillating between conservation and deep divergence. We found extensive sharing of the complements of receptor families expressed in neurons of the main olfactory epithelium (i.e., Olfrs, TAARs and MS4As) (Fig. 3A, Fig. S12, Fig. S13). Among *Peromyscus* Olfrs, 800 have homologs in both *Mus* and *Rattus* genomes, 302 of which are single-copy orthologs, 162 are either homologous to *Mus* or *Rattus* genes, and only 56 Olfrs are species-specific. Thus, despite having shared their last common ancestor 25 Mya and having experienced structural remodeling of their genomes, the house mouse, rat, and deer mouse do not differ markedly in the make-up of chemosensory receptor genes available to sample the chemical space covered by the main olfactory system.

**Fig. 3.**
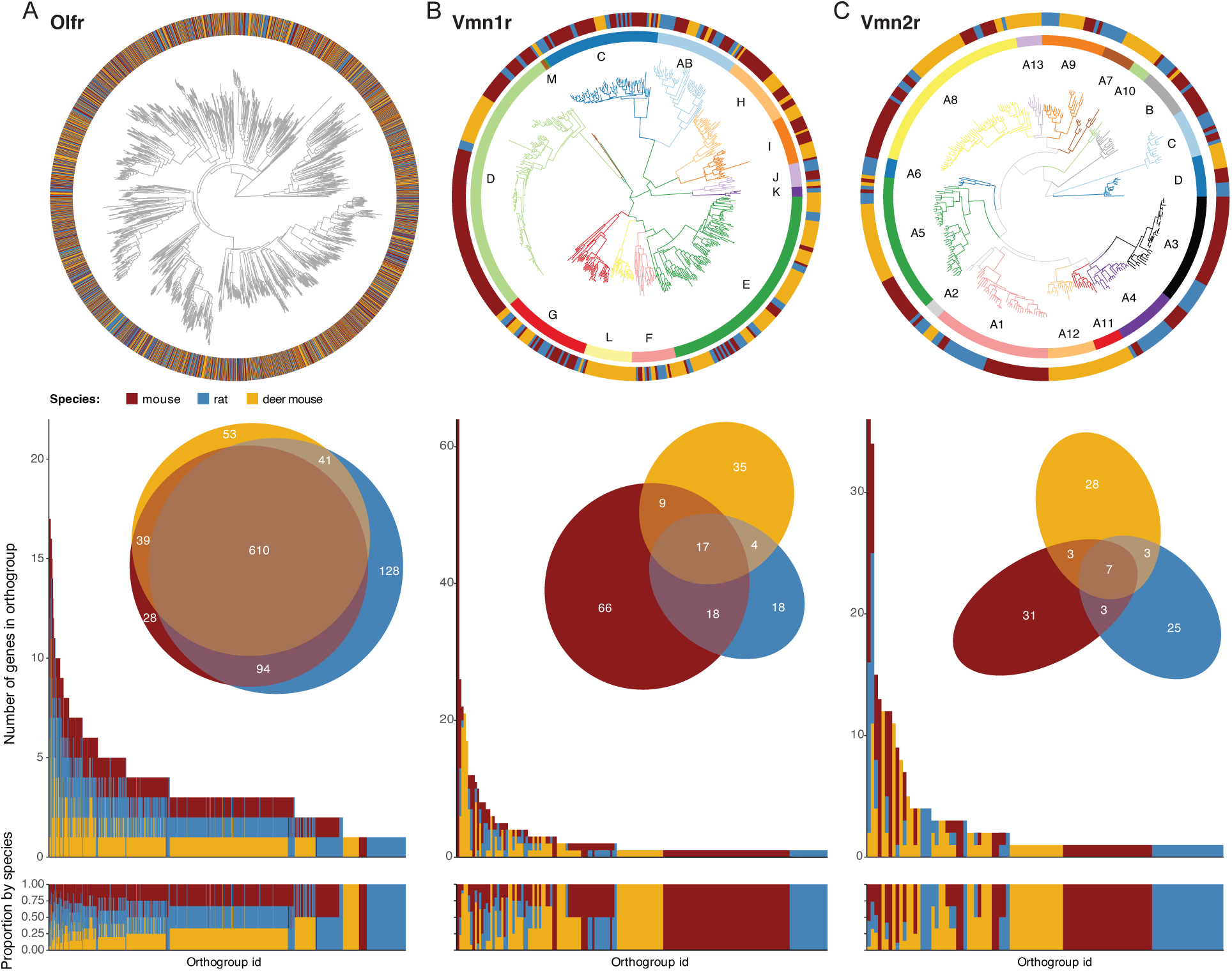
Comparison of the mouse, rat and deer mouse olfactory receptor gene repertoires. (**Upper row**) The phylogenetic gene trees depict the relationship among functional chemosensory receptor sequences. The outer rings are colored based on the species of the corresponding leaves on the tree. In the case of vomeronasal receptors (Vmn1r and Vmn2r), the families are indicated (color-coded inner circles). (**Lower row**) Genes were clustered into orthogroups on the basis of their amino acid sequence similarity. The y-axis of the top bar graph shows the number of genes of each species in each orthogroup, whereas it indicates the percentage of genes of each species within the orthogroup in the bottom bar graph. The x-axis of both top and bottom bar graphs indicates the orthogroup membership. The Venn diagrams show the number of orthogroups shared among species.

By contrast, we found that the receptor repertoires of the vomeronasal system differ noticeably: only 20% of Vmn1r and 7% of Vmn2r orthogroups are shared between the three taxa (Fig. 3B,3C). In the case of Fprs, Fpr-rs3 is the only member of the vomeronasal Fpr present in the deer mouse genome, which corroborates earlier reports that Fpr-rs4, -rs7, and -rs6 orthologs are found only in Murinae (*40*). At the level of vomeronasal receptors, we find that the clades previously identified are usually present but have experienced lineage-specific diversification resulting in ‘semi-private’ receptor repertoires. In some clades, there is evidence of diversification in *Peromyscus* (e.g., Vmn1rE and Vmn1rL), whereas others are reduced or absent (e.g., Vmn1rC, Vmn2rA3 and Vmn2rA4) (Fig. 3B,3C). One example of the latter includes *Vmn2r115* and *Vmn2r116* which encode receptors associated with the detection of the pheromone exocrine gland-secreting peptides (ESPs) in *Mus*, a class of tear secretion important for innate behavior in mouse (*41–43*).

### Conservation correlates positively with expression patterns

While cataloging the diversity of receptor genes among species provides insight into the drivers of selection and the functional requirements of each species (*32, 44–46*), estimating the abundance of sensory neurons expressing a given receptor provides complementary information to investigate the evolutionary dynamics of mammalian olfaction (*47*). Therefore, knowing the relative abundance of each receptor gene is a critical step towards a global understanding of the functional characteristics of the olfactory system. We thus investigated the gene expression profiles of olfactory tissues to better understand the evolutionary dynamics of these receptor families. Specifically, we performed analyses of bulk RNA-Seq data of the whole olfactory mucosa (WOM) and vomeronasal organ (VNO). Genome-wide expression profiles were highly correlated between the biological replicates for each sex and tissue of deer mouse samples (Fig. S14, S15), indicating strong overall similarity in intraspecific gene expression profiles, consistent with previous studies in laboratory (rat, mouse) and non-laboratory animals (dogs, marmosets, macaques, and humans) (*47–50*). Aside from the X-inactive specific transcript *Xist*, no genes exhibited significant sexual dimorphism in either deer mouse olfactory tissue, emphasizing the largely shared gene expression landscape between sexes, although we acknowledge that single cell sequencing might provide a more fine-grained analysis of subtle differences between the sexes. We found that nasal olfactory chemoreceptor genes are expressed across a broad dynamic range (Fig. S16), with a few highly abundant and most expressed at low levels. As receptor expression is positively correlated with the number of olfactory sensory neuron subtypes (*50*), these results indicate that the majority of sensory neuron subtypes are present in relatively low numbers.

To examine the relationship of sensory neuron subtypes across species, we first plotted the relative frequency of each subtype—defined as the receptor’s average expression represented as a percentage of the total average expression—onto the phylogenetic tree of each receptor family. We specifically focused on ORs and VRs, as their expression levels quantitatively reflect the abundance of sensory neuron subtypes and their sensitivity to receptor ligands (*6, 50*). Within each species, there is no apparent clustering of the genes that define the most abundant subtypes of sensory neurons (Fig. 4). In the Olfr tree, there appears to be a moderate level of conservation in OSN subtype across species (Fig. 4A). In contrast, we observe no apparent level of conservation in subtype representation in Vmn1r and Vmn2r trees (Fig. 4B,4C). To assess the degree of similarity in gene expression profiles between species, we calculated Kendall’s Tau correlation (τ) based on the summed percentage expression of genes within each orthogroup for each species. For Olfrs (Fig. 4D), The strongest correlation is between deer mouse and mouse (τ = 0.36), which indicates a moderate positive correlation and suggests that these two species share more similar expression patterns across orthogroups. There is less similarity between deer mouse and rat, which show a weak positive correlation (τ = 0.21), whereas mouse and rat exhibit a moderate positive correlation (τ = 0.35). Collectively, these data show a weak-to-moderate relationship in Olfr expression patterns, suggesting that there is mild conservation in the regulatory mechanisms affecting the abundance of OSN subtypes in the WOM. In the VNO, the near-zero correlations between species indicate negligible associations in gene expression profiles across species (Fig. 4E), which is expected given the limited overlap in gene repertoires. By contrast, gene markers of the major cell types show high level of expression similarities across the species analyzed (Fig. 4F,4G). Correlation analysis revealed that WOM marker gene expression profiles are highly conserved among species (τ = 0.70–0.78; Fig. 4F), suggesting strong evolutionary conservation. In contrast, VNO gene expression patterns exhibit moderate similarity across species (τ = 0.56–0.64; Fig. 4G), indicating both conservation and species-specific adaptations. Cross-tissue comparisons (WOM vs. VNO) showed weaker correlations (τ = 0.36–0.46), reflecting that their distinct functional roles are also mirrored in their molecular profiles (Fig. 4H).

**Fig. 4.**
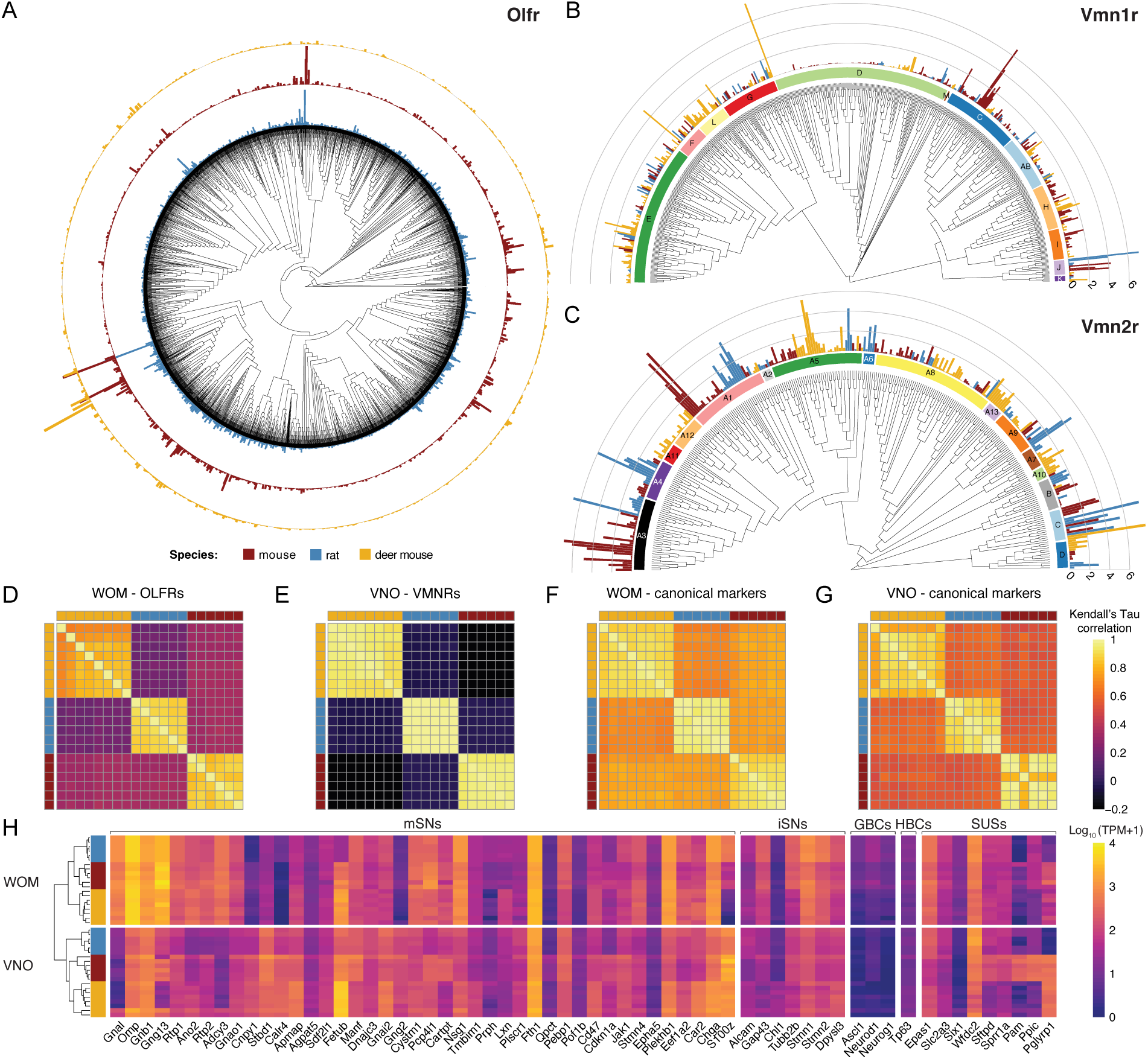
Comparative transcriptional landscapes of the olfactory and vomeronasal mucosa across species. Unrooted phylogenetic tree and mean expression levels for (**A**) Olfr, (**B**) Vmn1r, and (**C**) Vmn2r receptor families. Bars indicate the mean contribution (%) of each receptor to the total gene expression within each family and per species. Species color scheme used is kept consistent across all panels. Heatmaps of the pairwise Kendall’s Tau correlation coefficient among samples based on the expression levels of (**D**) OLFR, (**E**) VMNR orthogroups and canonical marker orthologs in (**F**) the WOM and (**G**) the VNO. (**H**) Heatmap of the expression levels of canonical markers for the main cell populations present in the whole olfactory mucosa (WOM) and vomeronasal organ (VNO). RNA expression levels of orthologs are represented on a log_10_ scale of transcript per million (TPM +1). mSNs: mature sensory neurons (OSNs and VSNs); iSNs: immature sensory neurons; GBCs: globose basal cells; HBCs: horizontal basal cells; SUSs sustentacular cells.

Taken together, these results indicate that the repertoires of chemoreceptors and their expression profiles have followed markedly different trajectories during the evolution of the olfactory sensory systems of rodents. In particular, given that the physiological responses of chemosensory neurons are largely determined by the chemoreceptor genes they express (*51*), our results support the idea that the functional tuning of the vomeronasal system evolves more rapidly than that of the main olfactory system. Furthermore, our data show that phylogenetic conservation correlates positively with receptor expression patterns.

### Conserved vomeronasal receptors exhibit conserved functions

Genes that exhibit relative stability over evolutionary time are more likely to encode essential functions, including the detection of ecologically relevant features. Our comparative genomic analyses reveal that a small set of vomeronasal receptor genes have been preserved in the *Mus* and *Peromyscus* genomes during evolution. This raises the question of the relationship between VR sequence and chemosensory function. Closely related receptors are likely to be activated by structurally-related ligands that may be associated with similar ethological contexts and even regulate similar types of behaviors. While the vast majority of vomeronasal receptors are orphan receptors with no known ligand, we identified a small set of receptors for which the cognate ligands have been identified in the laboratory mouse (Supplementary Table 2). These include several receptors for sulfated steroids and bile acids (BAs), both of which are components of urinary and fecal chemosignals that activate the accessory olfactory system (*52–56*)(Fig. 5A).

**Fig. 5.**
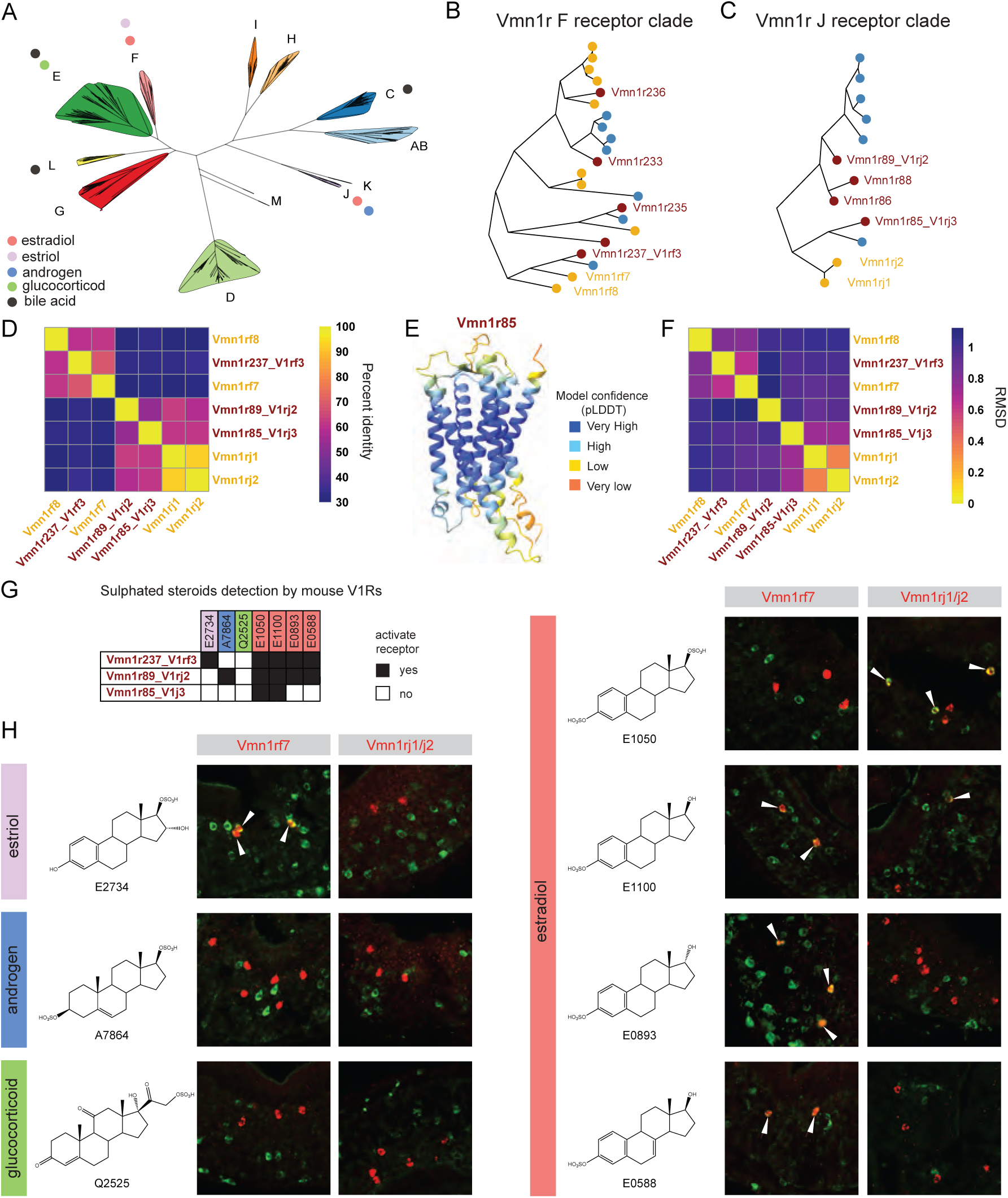
Detection of sulfated steroids by conserved V1Rs. (**A**) Mapping of cholesterol-derivatives detection on Vmn1r gene tree. Clades including specific receptors detecting each steroid class are indicated by dots. (**B**) Vmn1r F and (**C**) Vmn1r J receptor gene trees including homologs from mouse (crimson), rat (steel blue) and deer mouse (golden apricot). (**D**) Heatmap showing the percentage of protein identity for mouse Vmn1r237, Vmn1r89 and Vmn1r85 and their deer mouse nearest neighbor in the phylogeny. (**E**) Example of AlphaFold2 (AF2) predicted structure for a Vmn1r. The predicted confidence metrics (plDDT) indicate an overall high local accuracy in the models. (**F**) Comparison of AF2 predicted structures. For each pair, the Root Mean Square Deviation (RMSD) value is calculated based on pruned atom pairs in ChimeraX. (**G**) Summary of the detection of selected steroids by Vmn1r237, Vmn1r89 and Vmn1r85 in mouse. (**H**) Immunofluorescence with anti-pS6 (240/244) antibody (green) and in situ hybridization with RNA probe (red) after deer mouse VNO stimulation by individual steroids. Arrowheads mark colocalization between pS6 and receptor signals. The estriol E2734 only activates Vmn1rf7 whereas the androgen A7864 activates neither Vmn1rf7 nor Vmn1rj1/2. Vmn1rf7 is activated by most tested estradiols (E1100, E0893, E0588) but not by E1050. The estradiols E1050 and E1100 activate Vmn1rj1/2-expressing neurons. Overall, Vmn1rf7 in deer mouse shows a pattern highly similar to its mouse homolog Vmn1r237 (V1Rf3). Likewise, deer mouse Vmn1rj1/2 and mouse Vmn1r85 exhibit identical profiles.

The identification of putative cognate receptor-ligand pairs provided an opportunity to test whether conservation at the level of the primary sequences and proximity in terms of evolutionary distance are indicative of a high degree of functional similarity. To address this, we investigated the detection of sulfated steroids. Specifically, we tested the activation of the homologs of Vmn1r237 (V1rf3) and Vmn1r85 (V1rj3), which are respectively encoded by *Vmn1rf7* and *Vmn1rj1*/*Vmn1rj2* in *Peromyscus* as indicated by our orthology inference (Fig. 5B-C; Supplementary Table 2). The homologous proteins exhibit high to moderate primary sequence identity (Fig. 5D). To visualize and compare structural features between receptor homologs as well as their closest paralogs, we used AlphaFold 2.0 to generate models without a structural template (*57*). Overall, the monomeric models offer a reliable secondary structure, as suggested by the observed predicted confidence metrics (plDDT), and thus provide a first meaningful comparison of secondary structural elements and the general fold between vomeronasal receptors (Fig. 5E). Next, we generated pairwise superposition of receptor predicted structures to obtain root mean square deviations (RMSDs), a quantitative measure of similarity between two or more protein structures (Fig. 5F). The analyses indicate a high degree of similarity between the inferred homologs in comparison to their nearest paralogs, suggesting a possible correlation between structural features and degree of functional similarity. To assess the plausibility of this relationship, we tested the activity of neurons expressing those receptors when challenged with a selected set of ligands. The *Mus* receptors were previously shown to be activated by sulfated steroids (*52*). More specifically, Vmn1r237 and Vmn1r85 appear broadly selective of estradiols, estriols, but do not respond to androgens or glucocorticoids (Fig. 5G). In initial experiments, we assessed the induction of the phosphorylated ribosomal protein S6 (pS6), a sensitive molecular readout of VNO neural activity (*58*). Our data show that pS6 is a reliable marker of sensory activity following exposure to chemosignals in *Peromyscus* (Fig. S17). We then exposed animals to specific steroid ligands and used an *in situ* hybridization-based approach to identify the VRs expressed in pS6^+^ neurons identified using a pS6 240/244 antibody. Our data indicate that, when male deer mice were exposed to individual synthetic steroid sulfates, Vmn1rf7 was specifically activated by estradiols and estriol, but not androgens or glucocorticoids (Fig. 5H). The activity pattern we uncovered is reminiscent of the response of Vmn1r237 in *Mus* (*52, 54*). VNO neurons expressing *Vmn1rj1* and *Vmn1rj2* (denoted as *Vmn1rj1/j2* since they could not be distinguished by *in situ* hybridization) were activated by 17β-estradiol sulfate (E1050) and 17β-estradiol 3-sulfate (E1100). We did not notice any activation by estriol 17-sulfate (E2734), androgen (A7864) or glucocorticoid (Q2525) (Fig. 5H). The specificity exhibited by *Vmn1rj*-expressing neurons closely matches that of Vmn1r85 in *Mus* (*52–54*). We conclude that features of the primary sequences of these VRs are strongly correlated with their degree of functional similarity. Since excreted steroids are ubiquitous, though sensitive, reporters of an animal endocrine state and given the accumulated evidence of their importance in vertebrate social communication (*59*), our results identify a set of conserved V1Rs that act as detectors of the physiological status of an animal.

Among the V2R class of receptors, the ortholog of *Vmn2r65* in *Mus* is noteworthy as this gene encodes a receptor which is activated by the submandibular gland protein C (Smgc), a salivary protein produced by pup and adult female submaxillary salivary glands (*60*). In *Mus*, Smgc is a chemosignal involved in pup-directed behavior and knock-outs for *Vmn2r65* show a decrease in pup-directed attacks by normally infanticidal virgin males (*60*). Our data show that when male deer mice were exposed to pups, neurons expressing the ortholog of *Vmn2r65* were activated, indicating that this receptor may also be used to detect pup cues in *Peromyscus* (Fig. S14). While the identification of the precise ligand used by *Peromyscus* or its ethological significance are beyond the scope of this study, it is noteworthy that an ortholog of *Smgc* is present in the deer mouse genome.

Taken together, our results indicate that the relationships among receptor primary sequences are correlated with chemoreceptive similarity across species, revealing conserved receptor-ligand pairings within subsets of the vomeronasal sensory population.

## Discussion

We have generated a comprehensive whole-genome assembly and manually curated annotation –with a focus on the olfactory subgenome– for the North American deer mouse *Peromyscus maniculatus bairdii*. In part due to its well-documented natural history, the deer mouse represents a classic model system next to the house mouse for studying the genetic basis of adaptation at the morphological (e.g., coat color, tail morphology, sperm morphology), physiological (e.g., high altitude), and behavioral levels (e.g., parental behavior, burrowing behavior, nesting). Our high-quality chromosome-level genome assembly enables detailed gene-based studies, evolutionary analyses among populations, and facilitates transcriptomics and other genomics-based approaches in this model system. Indeed, this assembly has been adopted as representative genome by ENSEMBL and NCBI and has already empowered several studies since its public release (*39, 61–64*).

An annotated chromosome-level assembly further expands the ability to perform cross-species comparisons. We highlight the similarities and differences in genome organization compared to the murids *M. musculus* and *R. norvegicus.* In terms of gene content, we identified gene functions that show high volatility, revealing large differences between species. Specifically, genes involved in odor detection appear to be particularly susceptible to change, leading us to focus on chemosensory receptor genes. We show that variation introduced by large (i.e., intra- and inter-chromosomal rearrangements) and small-scale changes (i.e., tandem duplications) play major roles in generating volatility within the olfactory subgenome and ultimately the chemosensory gene repertoire.

Early genomic studies revealed pronounced differences in the chemoreceptor gene numbers and pseudogene proportions, a hallmark of the olfactory subgenome evolution (*12*). While rapid changes appear to be a signature of chemosensory receptor evolution, we highlight nuanced evolutionary trajectories among the gene families, ranging from a high degree of conservation in families associated with the main olfactory system to limited overlap in the receptor complements expressed in vomeronasal neurons. This likely results from differences in selective pressure, a major factor determining how evolution shapes the trajectories followed by the different chemosensory gene families (*2, 65*). With genome rearrangements common in mammals, and particularly in rodent genomes as illustrated in our analyses, there are theoretically ample opportunities for gene loss around breakpoints. Olfrs constitute the largest class of membrane receptors in mouse and are typically found in large clusters distributed across the genome (*48*). Olfrs are characterized by a high degree of functional redundancy and their ligands are low molecular weight volatile molecules. Functional redundancy would suggest that some genes can likely be lost without dramatic consequences, yet our results argue that events causing the loss or disruption of OR clusters likely pose selective disadvantages and are disfavored evolutionarily. Consequently, we observe a high retention of homologous olfactory receptor genes, indicating extensive functional overlap among the surveyed species in the detection of environmental odors (e.g., food, habitat). Besides its role in detecting odors from the chemical environment of an animal, the main olfactory system is also essential for proper social behavior, as mice without a functional MOE exhibit deficits in sexual and defensive behaviors (*66*), therefore adding additional selective constraints. Consistent with this, we observed the conservation of ORs detecting semiochemicals such as *Olfr1509* (activated by (methylthio)methanethiol (MTMT), a strong-smelling component of male mouse urine that attracts female mice (*67*))*, Olfr1019* and *Olfr1395* (activated by 2,4,5-trimethyl thiazoline (TMT), a red fox odor that elicits aversion and fear (*68, 69*)). Purifying selection has thus promoted the evolutionary conservation of a significant swathe of genes encoding chemosensory receptors expressed by neurons of the main olfactory epithelium.

In sharp contrast, vomeronasal receptor repertoires appear highly divergent, likely reflecting the necessity to detect more dissimilar sets of ligands. One possible explanation for the distinct retention patterns of VNO versus MOE receptor genes lie in the information conveyed by their ligands. Historically, the vomeronasal system was considered pheromone-specific (*70, 71*), with the rapid evolution of the VR gene repertoire proposed to reflect the species-specific nature of pheromones and the need to respond to very different ligands (*72, 73*). However, recent findings in mouse indicate that many vomeronasal sensory neurons are activated by non-pheromonal ligands (*74*) and the majority of VRs recognize heterospecific cues (*52*). The two classes of vomeronasal receptors –Vmn1r and Vmn2r– use distinct strategies to encode chemosensory information, and distinct receptor subfamilies have evolved towards the specific recognition of certain animal groups or chemical structures (*52*). Vmn2rs appear tuned to cues associated with the identity of the emitters and with clear behavioral significance, such as the sex of a conspecific or the predatorial or competitive nature of a heterospecific, whereas Vmn1rs appear to encode other types of biologically relevant information related to the physiological status of the emitter (*52*).

While ligands for the vast majority of VRs are unknown, the limited list of receptor-ligand pairs allows to confirm general principles regarding the type of cues each class of VRs is tuned to: Vmn1rs are dedicated to the detection of small molecules typically present in excretions of mammals and other animals (e.g., fecal bile acids (*55, 56*), sulfated steroids (*52, 54*)), whereas Vmn2rs are associated with the detection of peptides and proteins (e.g., exocrine-gland secreting peptides 1 and 22 (ESP1, ESP22) (*41, 75, 76*), cystatin-related protein 1 (CRP1) (*77*), hemoglobin (*60, 78*), and urinary proteins (MUPs) (*74, 79*). Thus, the odor space covered by Vmn1rs vs Vmn2rs appears to separate metabolites from gene products. Metabolites are produced by more or less conserved pathways across mammals and vertebrates, such as the excretion of steroid derivatives or bile acids. Importantly, besides host molecules, it is likely that microbial metabolites derived from bacterial metabolism of dietary substrates, the modification of host molecules such as conjugated bile acids or produced directly by the gut microbiota are also ligands of VNO neurons. With 20% of Vmn1r orthogroups conserved across taxa, our results indicate that some V1R receptors tuned to detect these ligands are evolutionarily conserved. On the other hand, the ligands for V2R receptors, excreted peptides and proteins, are a direct reflection of the encoding genes and appear more divergent in nature. The few characterized ligands appear to undergo rapid molecular evolution. For example, the expansion of exocrine-gland secreting peptides (ESPs) seems to be substantial in murids, with the ESP family consisting of 38 members in mice and 10 members in rat (*75, 80*) and few to no representatives in the deer mouse genome (this study). Similarly, proteins of the major urinary protein (MUP) complex are diverse in mouse and rat (*79, 81*), but missing in *Peromyscus*. These expansions in *Mus* correlate with co-expansion of Vmn2rs (*79*). Conversely, we see the opposite pattern in *Peromyscus*, with co-contraction in the corresponding receptor clades and the volatility of peripheral circuit elements involved in the perception of social cues.

Our integrative approach has permitted both the identification of candidate genes involved in rapidly diverging traits between species and evolutionarily stable genes encoding essential chemosensory functions, foremost the detection of biologically relevant features. Deer mice are one of the best studied rodents in terms of behavior and ecology and the chromosome-level genome assembly reported here has already and will continue to serve as an important resource to further its role as a model clade (*18*). The receptor catalogue uncovered in this study provides a robust molecular foundation to further dissect the neural circuits governing chemosensation in rodents and define how neural circuits evolve in response to the rapid changes in receptor repertoires of the peripheral nervous system.

## Materials and Methods

### Animal husbandry and ethical statement

We established colonies of *P. maniculatus bairdii* (strain BW) at Harvard University and the University of Liège from animals originally obtained from the Peromyscus Stock Center at the University of South Carolina. We housed animals in barrier, specific-pathogen-free conditions with 16 h light: 8 h dark at 22 °C in individually ventilated cages. Animals were provided irradiated mouse diet *ad libitum* and had free access to water. We weaned animals at 23 days of age into cages with at most five members of the same sex. After weaning, we fed animals irradiated diet *ad libitum* with free access to water and provided them with nesting material and a polycarbonate translucent red hut or tube. Animal procedures were conducted with approval by the Institutional Animal Care and Use Committees (IACUCs) of Harvard University (protocol 13-03-148) and the University of Liège (protocol 22-2449).

### Genome Sequencing and Assembly

To improve the quality and contiguity of a previous deer mouse *P. maniculatus bairdii* genome assembly (Pman1.0; GCF_000500345.1) we sequenced reads from long-jump *Fosill* libraries and produced a new assembly by combining those reads with the publicly-available short-read data produced to generate the first draft assembly. Specifically, DNA was extracted from the liver of a single female individual obtained from our laboratory colony (BW stock) using the Qiagen Blood & Tissue kit following the manufacturers’ instructions and used as template to produce long-jump *Fosill* libraries as described in (*82*). The choice of a female individual means that equal coverage is obtained for the autosomes and the X chromosome, while the Y chromosome is not included in the final assembly. *De novo* sequencing of *the Fosill* libraries was performed on an Illumina HiSeq 2500 at the Broad Institute Sequencing Platform (Cambridge, MA, USA). All data from short paired-end libraries and longer mate-pair libraries were assembled using the ALLPATHS-LG genome assembler v.R4855 (*83*) (Supplementary table 1). Details about the assembly statistics are given in Supplementary table 2.

To anchor scaffolds into chromosomes, we took advantage of genetic linkage data to determine the placement, ordering, and relative orientation of scaffolds into each chromosome. We used ALLMAPS v.0.6.2 (*84*) to combine the information from several published maps generated using interspecific crosses between *P. maniculatus bairdii* (BW stock) and *P. polionotus subgriseus* (PO stock)(*19–21, 24*) and genetic information from an intraspecific cross between *P. maniculatus bairdii* (BW) and *P. maniculatus nubiterrae* (NUB)(*61*). Most of these genetic linkage maps were constructed using ddRAD-seq (double digest restriction site-associated DNA sequencing (*85*)). We gathered the consensus sequences corresponding to the markers and aligned them to the genome scaffolds using BLAT v.36 × 2 (*86*). We filtered markers to keep only those that had a unique match in the genome. The final dataset comprised 63,318 ddRAD markers plus 188 gene-based markers included in the genetic linkage map constructed by Kenney-Hunt *et al.* (2014)(*24*). During a first iteration, ALLMAPS revealed that a total of seventeen scaffolds contained markers associated with more than one linkage group. These were likely misassembled and split accordingly. The breakpoints positions were determined based on the ALLMAPS prediction and the detection of discordantly mapped reads. In most cases, these corresponded to assembly gaps.

Our final assembly includes 585 scaffolds, encompassing 2,436,387,942 bp (97% of the total sequenced bases), distributed in 23 autosomes plus the X chromosome. The orientation of 447 scaffolds, corresponding to 2,427,689,157 bp (96.6% of the total sequence), could be determined due to the presence of more than one marker. We assigned chromosome names based on previous reports from interspecific reciprocal whole chromosome painting, which have allowed to assign linkage groups with known genes to *Peromyscus* chromosomes (*24, 25*). The chromosome names reported here reflect the standardized *Peromyscus* cytogenetic nomenclature (*87*).

### Repetitive sequences identification and enrichment analyses

Repetitive sequences were annotated using a combination of homology searching and *ab initio* prediction as described previously (*39*). To test for an enrichment of repeats in regions harboring chemosensory genes, we performed a permutation test for enrichment of repeats for five types of repeats. Specifically, we focused on three classes of long terminal repeats (LTR), namely endogenous retroviruses (ERVs) of class I (ERV1), class II (ERVK) and class III (ERVL), and two classes of non-LTR retrotransposons, namely long interspersed nuclear elements (LINE) and short interspersed nuclear elements (SINE). We extracted the flanking regions of each gene using bedtools flank (version 2.29.1) and compared repeat composition for these regions relative to expectations from randomization across 1,000 permutations using GAT (version 1.3.5) (*88*).

### Whole genome alignment and comparative annotation

To annotate the genome, we used an annotation strategy making use of multiple genome alignments and an existing high-quality annotation set (*28*). We first aligned the deer mouse chromosome-level assembly to the assemblies of the house mouse (*Mus musculus;* GRCm38), the rat (*Rattus norvegicus*; Rnor_6.0), the prairie vole (*Microtus ochrogaster;* MicOch1.0) and the oldfield mouse (*Peromyscus polionotus;* Ppol.1.3) using ProgressiveCactus v.0.0 (*89, 90*). We reasoned that including more species that represent progressive levels of evolutionary divergence would improve the accuracy of the ancestral sequence reconstruction process that takes place during the preparation of the whole-genome alignment. Using the Comparative Annotation Toolkit (CAT v.4c4140; (*28*)), we annotated the deer mouse genome using the genome of *Mus musculus* (GRCm38/mm10) and the high-quality and well-curated GENCODE VM15 as the reference gene/transcript set, as well as extensive RNA-Seq datasets for *P. maniculatus bairdii* derived from six tissues (brain, testis, hypothalamus, skin, whole olfactory mucosa and vomeronasal organ).

To annotate genes encoding chemosensory receptors, we first created a customized file containing all annotated ORs, V1Rs and V2Rs for mouse, rat and deer mouse (from *de novo* transcriptome assemblies and RefSeq from Pman1.0). We mapped the protein sequences against the deer mouse genome using Spaln (*91*). In parallel, we aligned trimmed-reads against the genome using HISAT2 version 2.2.0 (*92*) and used StringTie version 2.1.3b (*93*) with the RNA-Seq alignments to reconstruct a non-redundant set of transcripts observed in any of the RNA-Seq samples. The diverse evidence sources were loaded in an Apollo workspace (*94*) which was used to perform manual curation. The Apollo and CAT annotations were then merged using gff3_merge tool from the GFF3 Toolkit (*95*). Finally, we applied Trinotate version 3.2.1 (*96*) to generate a functional annotation of the predicted proteome.

### Evaluation of genome completeness

To obtain quantitative measures of the completeness of the genome assembly, we used BUSCO (version 3.0.2) with BLAST+ (version 2.2.28+), HMMER (version 3.1b2) and AUGUSTUS (version 3.3.2). We used *human* as species, which specifies the parameters used by AUGUSTUS, and the *mammalia* and *euarchontoglires* sets for our analyses.

### Conserved synteny and riparian plot

To compare and visualize conserved synteny of orthogroups across *Rattus norvegicus*, *Mus musculus*, and our *Peromyscus maniculatus* annotation and assembly, we used the R package GENESPACE v1.3.1. For Rnor, we downloaded the translated coding sequences (CDs) and corresponding annotation file from NCBI (Rnor 6.0). For *Mmus* and *Pman*, we used AGAT v1.2.0 to generate translated CDS fasta files based on Gencode vM15 / GRCm38.p5 and our in-house generated annotation and assembly, respectively. Based on these input files, orthogroups and syntenic blocks were determined by GENESPACE’s main pipeline, which internally runs OrthoFinder and MCScanX. We visualized the results using GENESPACE’s *plot_riparian()* function using Pman as the reference genome and setting the useOrder=FALSE flag to order and scale the riparian plot by physical position.

To assess structural variation across *Rattus norvegicus*, *Mus musculus*, and *Peromyscus maniculatus* genomes we aligned them using the NUCmer module of the MUMmer package v3.23 (*97*). Mummer alignment plots and chromosome painting plots were generated using R.

### Functional enrichment analyses

For functional enrichment analyses, we used the WEB-based Gene SeTAnaLysis Toolkit (WebGestalt: https://www.webgestalt.org) (*98*). We performed an over-representation analysis (ORA) using the gene ontology (GO) database for biological process with *Mus musculus* as reference organism and genome protein-coding as reference set, and defining the list of protein-coding genes not transferred to the deer mouse genome during the comparative annotation as gene list. We used the default parameters and applied an adjustment for multiple comparisons based on a *P*_FDR_ < 0.05 (Benjamini–Hochberg method).

### Gene family evolution and positive selection

Gene family expansion and contraction analyses were performed using CAFÉ (v5.0) (*34*). Predicted proteomes were obtained from ENSEMBL and we used orthogroup counts derived from OrthoFinder version 2.5.5 (*99*) clustering of protein-coding genes from *Peromyscus maniculatus*, *Mus musculus, Rattus norvegicus*, and 14 mammalian species. Using the species tree calculated by OrthoFinder (*100, 101*), we generated an ultrametric species tree using the *make_ultrametric.py* script from the OrthoFinder distribution and calibrated it with molecular time estimates for divergence between pig (*Sus scrufa*) and mouse (*Mus musculus*) obtained from TimeTree (http://www.timetree.org). Error models accounting for potential biases in genome annotations were calculated using the built-in error correction procedure in CAFÉ. Statistical significance of gene family size changes along branches was assessed at p-values <0.05 using 1 to 5 Gamma categories with Poisson distribution for the root frequency distribution and with gene families that do not exist at the root. We compared and ranked models using the Akaike Information Criterion (AIC), calculated using the log-likelihood (lnL) from the CAFÉ output files. Gene ontology terms were assigned to an orthogroup based on the annotations of its members. We retrieved GO terms from the Trinotate annotation for deer mouse and ENSEMBL annotations for mouse, rat, vole and Chinese hamster. A given GO term had to be harbored by at least two members of the orthogroups to be considered representative of the family.

### Phylogenetic analyses of chemosensory genes

To investigate the evolutionary relationship of the deer mouse chemosensory genes, we carried out phylogenetic analyses with other rodent receptor proteins. Our focus was on mouse and rat, two species for which most is known in terms of receptor function. Protein sequences for these receptors were downloaded from GenBank. Protein sequences for *P. maniculatus* were predicted from the genome with our curated annotation. We aligned amino acid sequences using MAFFT (*102*) using the L-INS-i procedure and the BLOSUM45 matrix as the scoring matrix. The phylogenetic trees were constructed in IQ-TREE 1.6.11 (*103*). An automatic model search was performed using ModelFInder (*104*) with the search restricted to models including the WAG, LG, and JTT substitution models. The maximum likelihood analysis was performed using default settings and by calculating Shimodaira-Hasegawa-like approximate ratio test (SH-aLRT) support and ultrafast bootstrap support (UFBoot (*105*)) after 1000 replicates each. We used the R package *ggtree* (*106*) for visualizing and annotating the phylogenetic trees, with midpoint rooting performed using the midpoint.root function implemented in the *phytools* package (*107*).

### Comparative analyses of chemosensory gene repertoires

To compare chemosensory gene repertoires among species, orthogroups for *M. musculus*, *R. norvegicus*, and *P. maniculatus* were reconstructed using OrthoFinder. Analyses were performed independently for each chemosensory gene family (namely, Olfr, Ms4a, Taar, Vmn1r, Vmn2r, and Fpr). Gene counts, including singletons not assigned to any orthogroup, were imported into R for visualization using the *tidyverse* (v1.3.0) and *eulerr* (v6.1.0) packages.

### RNA-Seq Library Preparation and Sequencing

To estimate mRNA abundance in the whole olfactory mucosa (WOM) and vomeronasal organ (VNO), we conducted an RNA sequencing (RNA-Seq) experiment. Specifically, we euthanized four male and four female by CO_2_ inhalation for 2 min and collected the tissues. We immersed the tissue in ice-cold TRIzol reagent (Thermo Fisher, Waltham, MA, USA) and disrupted the tissues with a motorized pestle before storage at −80 °C until processing. Total RNA was extracted using Direct-zol-96 RNA (Zymo Research, Irvine, California, USA) and used as input to isolate messenger RNA with a NEB magnetic mRNA isolation kit (New England Biolabs, Ipswich, MA). We constructed complementary DNA (cDNA) libraries using the NEBNext Ultra Directional RNA Library Prep Kit for Illumina (NEB, Ipswich, MA). Multiplexed libraries were sequenced on an Illumina HiSeq 2500 (Illumina, San Diego, CA) with HiSeq version 4 chemistry in high output mode and paired-ends (2 × 125 bp). Libraries from males and females from both species were sequenced in the same lanes to alleviate batch effects. The average depth per sample was 28 million read pairs.

### RNA-Seq Data Analysis

In addition to data generated as part of this study, we retrieved RNA-Seq datasets available in the Sequence Read Archive (SRA) (study accession numbers: mouse: PRJEB1365 and PRJEB2572; rat: PRJEB1924). Raw sequencing reads were quality-controlled using FastQC and adapters/low-quality bases (quality score < 20) removed using cutadapt v.2.3 (*108*) while applying a minimum length cutoff of 36 bp. Trimmed reads were pseudoaligned against the corresponding transcriptomes using Kallisto (*109*). Transcriptome for deer mouse for extracted from our in-house genome annotation using gffread (*110*), whereas the annotations for mouse (*Mus musculus*; GRCm39) and rat (*Rattus norvegicus;* mRatBN7.2) were downloaded from NCBI (https://ftp.ncbi.nlm.nih.gov/genomes/refseq/vertebrate_mammalian). Transcript-level estimates and read counts were imported to R using the *tximport* package (*111*) to obtain length-scaled gene-level transcripts per million (TPM) values. We used DESeq2 (*112*) for count-based differential expression analyses in deer mouse using the negative binomial likelihood ratio test with an adjusted P value of P < 0.05.

### Sulfated steroid and stimulus exposure

Steroids were purchased from Steraloids Inc. (Newport, RI, USA) and diluted in 50% methanol (in water) to prepare 10 mM stock solutions stored at -20°C. Individual steroids were used at 500 µM in phosphate-buffered saline (PBS). 5 µL of steroid solution were spotted on each nostril of male BW animals (8-16 weeks), and the animals were exposed to steroids for 60 min. Experiments were conducted for a least two animals.

For the exposure experiments involving bedding stimuli or pup, a subject deer mouse was introduced (male BW; 8-16 weeks) into a clean cage for habituation. Upon addition of the test odor source, the individual voluntarily made direct contact with the stimuli in freely behaving conditions. The animals were exposed for 55 to 60 min.

Following exposure, the animals were sacrificed using a lethal injection of sodium pentobarbital (Euthasol) administered intraperitoneally and the VNO was promptly dissected in cold 1× PBS under a microscope. The dissected VNO were then immersed into freshly prepared 4% paraformaldehyde (PFA; Electron Microscopy Sciences) in PBS for 10 min on ice. Following fixation, the tissue was briefly rinsed in 1× PBS, transferred in a 30% sucrose solution in PBS on ice until it sank, and embedded in a tissue-freezing medium (OCT; Tissue-Tek) before freezing on dry ice. Samples were stored at -80°C until cryosectioning. We sectioned VNO coronally at 16 µm using a cryostat at -20°C (Leica CM3050 S, Leica Biosystems Inc., Buffalo Grove, IL) and collected the tissue sections on SuperFrost Plus Slides.

### In Situ Hybridization with pS6 immunostaining

To prepare RNA probes, we cloned complementary DNA of approximately 900-base-pair (bp) fragments into pCRII-TOPO (Invitrogen). Antisense cRNA probes were synthesized by in vitro transcription from PCR-amplified templates using T7 or Sp6 RNA polymerase (Promega). Digoxigenin (DIG)-labeled probes were used for receptor detection, and fluorescein (FITC)-labeled probes were used specifically for *Gnai2*, with labelling mixes sourced from Roche.

For single or dual color fluorescence *in situ* hybridization, we used labelled cRNA probes at 2 ng/µL and used a hybridization temperature of 68°C. First, the tissue sections were fixed in 4% PFA/1× PBS for 10 min and washed 3 times with 1× PBS for 3 min each. The sections were treated with acetylation solution (0.1 M triethanolamine with 2.5 μL/mL acetic anhydride) for 10 min. After 3 washes with 1× PBS, each for 5 min, the slides were incubated with the pre-hybridization solution (50% formamide, 5× SSC, 5× Denhardt’s, 2.5 mg/mL yeast RNA, 0.5 mg/mL herring sperm DNA) for 2 h at room temperature. Hybridization buffer was prepared by adding 4% dextran sulfate (Millipore) to pre-hybridization buffer and labelled probes (1 µL of each probe. The mixture was heated at 80°C for 5 min and immediately chilled on ice for 5 min. 125 µL of hybridization solution was applied to the slides, which were covered with HybriSlip hybridization covers (Invitrogen) and incubated in a sealed chamber for 16 h at 68°C. Following hybridization, the slides were washed with 5× SSC once for 5 min, and with 0.2× SSC three times, each for 20 min at 68 °C. Slides were washed at room temperature with 0.2× SSC for 5 min and subsequently with TNT buffer (100 mM Tris, pH 7.5, 150 mM NaCl, 0.05% Tween 20) for 5 min.

For single channel fluorescence, after the post-hybridization washes, 200 µL of anti-DIG-POD solution (Roche; 1/250 in TNB blocking buffer (Perkin-Elmer)) was applied to the slides which were covered with parafilm and incubated in a sealed chamber for 3 h at room temperature. The slides were then washed with 6 times with TNT buffer for 10 min each. The signal was developed for 10 min using the TSA Plus Cyanine 3 kit (Perkin Elmer) as per manufacturer’s protocol. Slides were washed with TNT (3 times, 5 min each and once for 1 h) and then transferred into PBS containing 0.05% Tween 20 detergent (PBST). 500 µL of blocking buffer containing 1× PBS / 5 % normal goat serum /0.1 % Tween 20 was applied on the slides for 1 h at room temperature. pS6 antibody solution (anti-phosphorylated ribosomal protein S6 rabbit polyclonal (Ser240/244) Antibody, Cell Signaling #2215, at 1/250 dilution in antibody dilution buffer containing 1×PBS/ 1%BSA/0.1 % Tween 20) was applied on the slides, which were covered with parafilm and incubated in a sealed chamber for 40 h at 4°C. After washing the slides 4 times 5 min with PBST, signals were visualized with Alexa647-conjugated goat anti-rabbit secondary antibody (Invitrogen #A21245) at 1:250 in antibody dilution buffer incubated overnight at 4°C. After washing 4 times 5 min at room temperature with PBST buffer, the slides were treated for 10 min with PBS containing DAPI (2 µL in 50 mL; 1 μg/mL DAPI), rinsed twice for 3 min with PBS and mounted with cover glass with Fluoromount-G (Invitrogen). Microscopy images were acquired using an Axio Imager Z2 (Zeiss).

Dual color fluorescence *in situ* hybridization combined with immunostaining was conducted as above with the following modifications. After the post-hybridization washes, 200 μL of anti-FITC-POD (Roche, at 1/250 dilution in TNB blocking buffer, Perkin-Elmer) was applied and incubated for 3 h at room temperature. After washing the slides, signal was developed for 10 min using the TSA Plus Fluorescein kit (Perkin Elmer). The slides were washed with TNT buffer 3 times, each for 5 min, and subsequently treated with 3% H_2_O_2_/1× PBS to suppress residual peroxidase activity. Slides were washed again 3 times with 1× PBS and TNT, each for 5 min. DIG antibody solution was applied to the slides which were covered with parafilm and incubated in a sealed chamber overnight at 4°C. Washes, TSA development, immunostaining was performed as described above.

### Protein Structure Prediction

Monomer structures of vomeronasal receptors were predicted using AlphaFold2 version 2.3.1 (*57*) run locally without using structural templates. Structures were visualized and analyzed in ChimeraX v 1.9 (*113*). Structural similarity between monomer models was quantitatively assessed using Root Mean Square Deviation (RMSD) calculations following pairwise structural superposition of predicted receptor models matchmaker tool within ChimeraX with default parameters for chain pairing (i.e., best-aligning pair of chains between reference and match structure), alignment (i.e., Needleman-Wunsch sequence alignment algorithm and BLOSUM-62 matrix), and fitting (i.e., iteration by pruning long atom pairs with an iteration cutoff distance of 2.0 Å).

## Supporting information

Complete set of CAFE_gene_families_with functions

Complete set of orthogroups with conserved receptors with annotations

## Acknowledgments

The computations in this paper were run on the FASRC Odyssey and Cannon clusters supported by the FAS Division of Science Research Computing Group at Harvard University.

## Author contributions

Conceptualization: JML, HEH. Formal analysis: JML, AFK, LG. Investigation: JML. Writing—original draft: JML. Writing—review & editing: JML with input from all authors. Visualization: JML, AFK, LG. Supervision: HEH.

## Data and materials availability

The *P. maniculatus bairdii* reference genome and the associated sequencing data are archived in the NCBI Datasets (BioProject no. PRJNA494228). Scripts and pipelines used for genome assembly, annotation, expression analysis, and phylogenetic analyses are available upon reasonable request to the corresponding author.

## Supplementary Materials

**Fig. S1.**
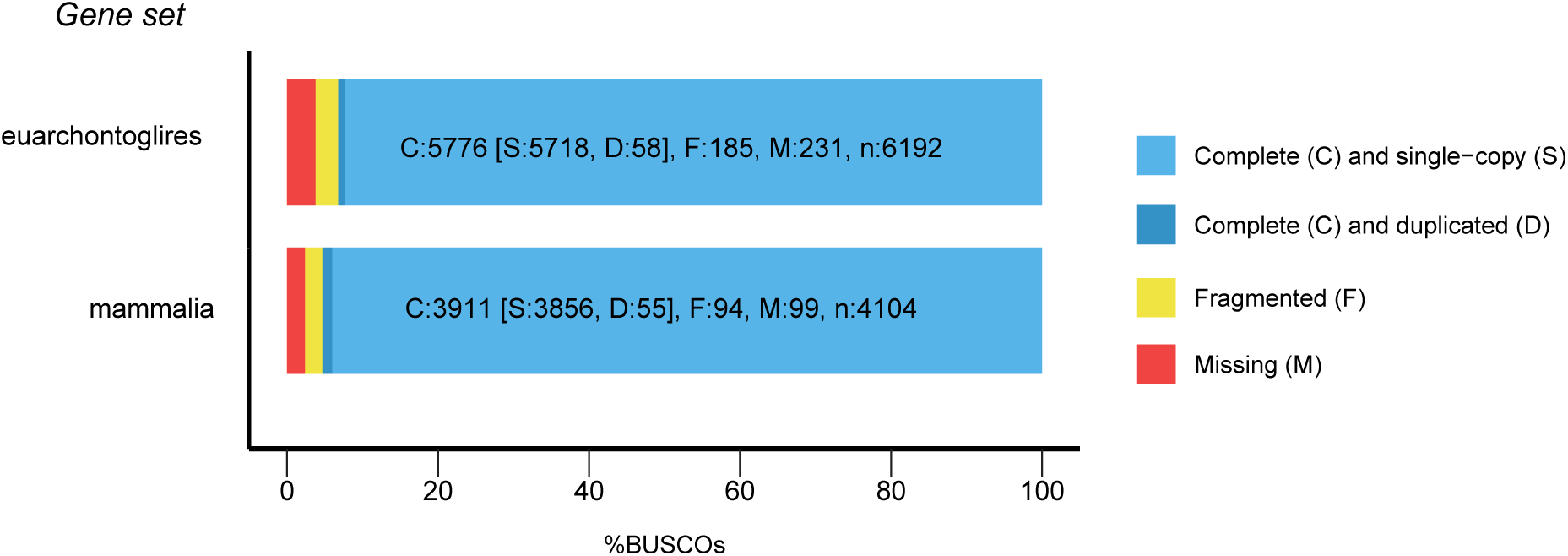
Summary assessments of genome completeness. The bar charts for the proportion of single-copy orthologs present in the HU_Pman2.1.3 version of the deer mouse genome inferred from search against two databases (BUSCO euarchontoglires_odb9 and mammalia_odb9 databases). Light blue shows the percent of complete and single copy genes, dark blue shows percent complete and duplicated genes, yellow shows the percentage of fragmented genes and finally red shows the % of missing genes in the assembly.

**Fig. S2.**
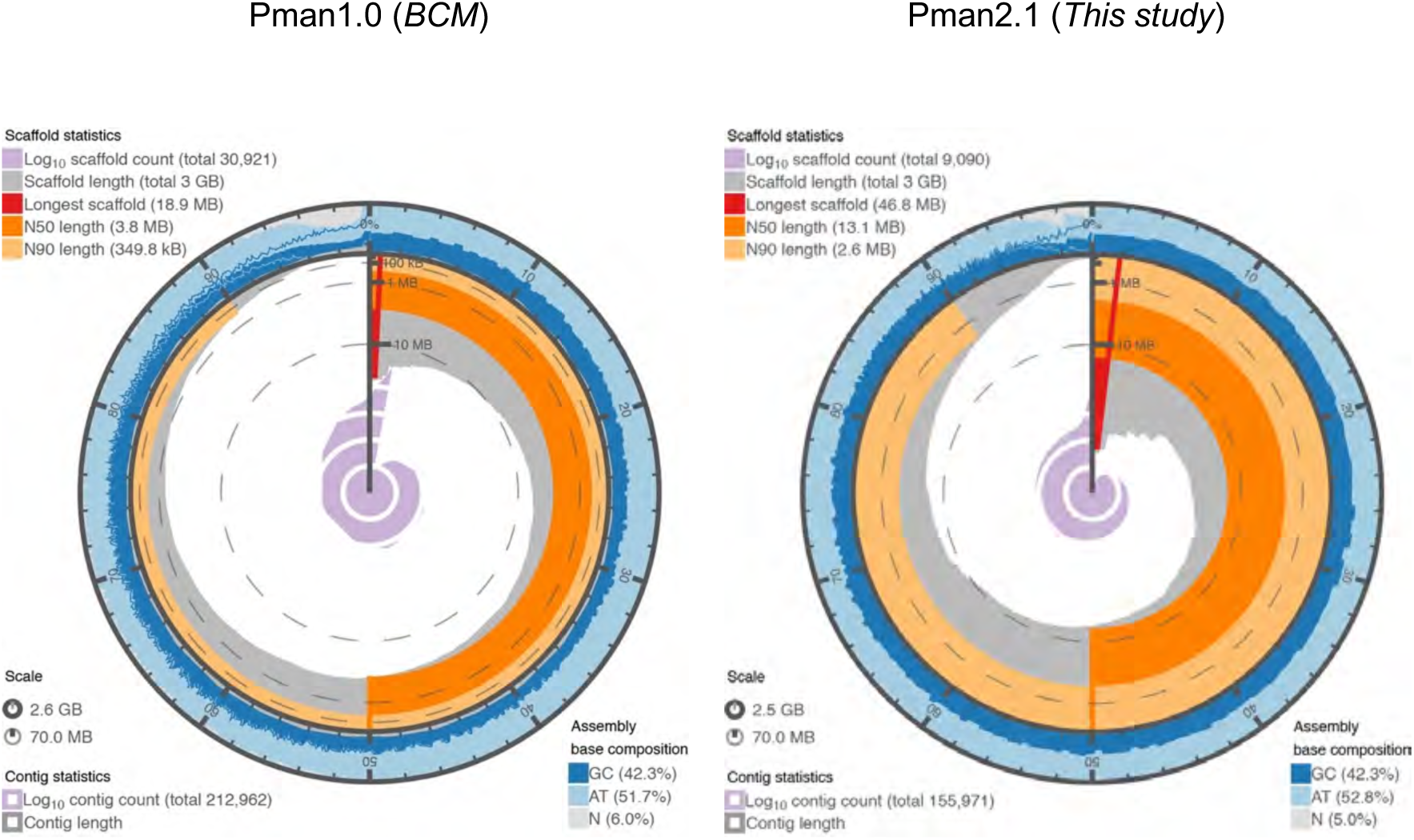
Comparison of assembly statistics for two versions of the *P. maniculatus bairdii* genome. The addition of data from fosmid jump libraries lead to an increase in the scaffold N50 and N90 values, a decrease in the total number of contigs and scaffolds, and an overall improvement of the quality of the raw assembly.

**Fig. S3.**
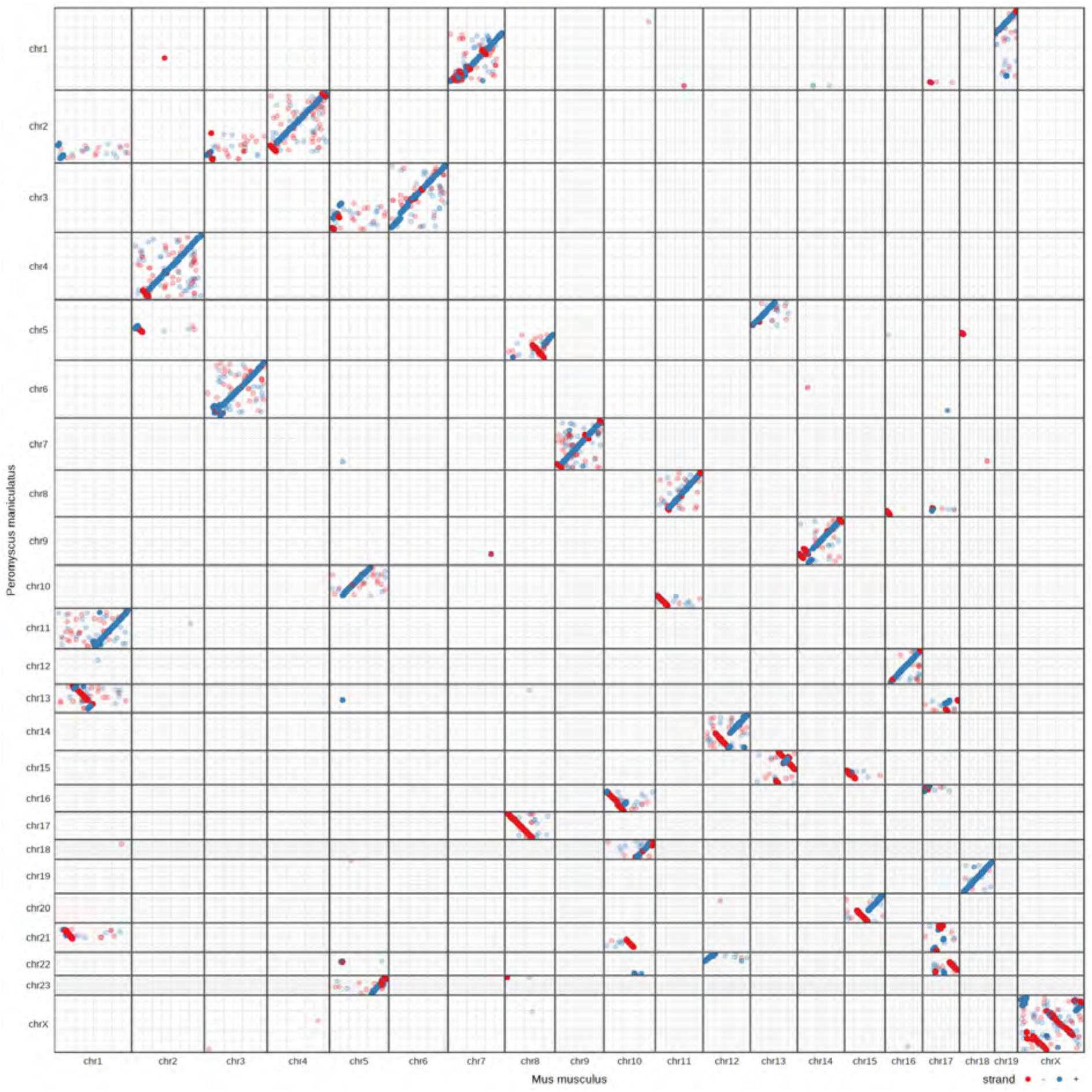
Dotplot of syntenic blocks between the laboratory mouse *Mus musculus* and the North American deer mouse *Peromyscus maniculatus* chromosomes. The blue and red lines indicate forward and reverse syntenic relationships, respectively.

**Fig. S4.**
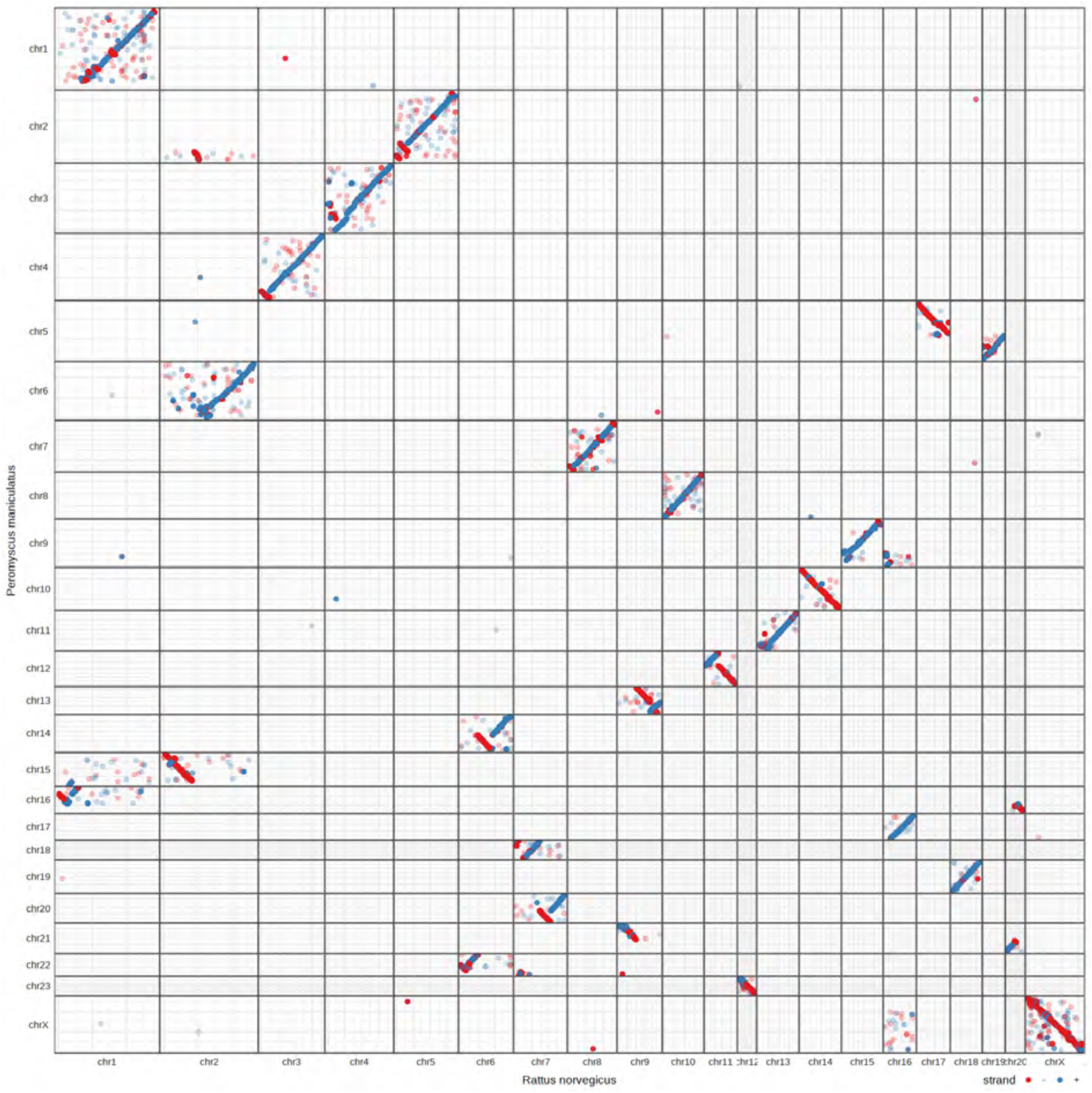
Dotplot of syntenic blocks between the Norway rat *Rattus norvegicus* and the North American deer mouse *Peromyscus maniculatus* chromosomes. The blue and red lines indicate forward and reverse syntenic relationships, respectively.

**Fig. S5.**
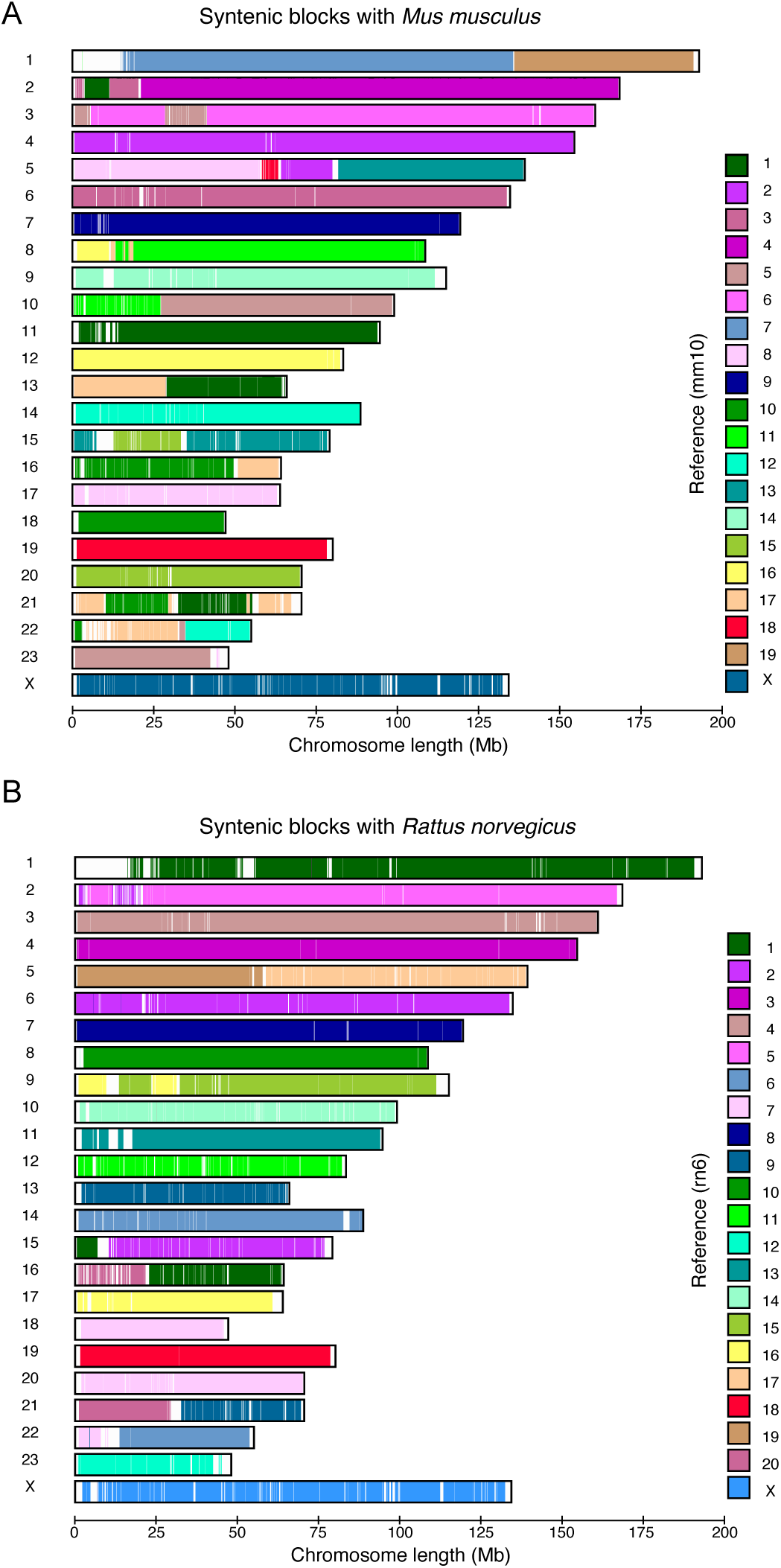
Ideogram of deer mouse chromosomes. Rectangles represent chromosomes, and colored blocks inside represent regions of homology with mouse (**A**) or rat (**B**) chromosomes. White filling represents regions of the deer mouse genome without homologous regions.

**Fig. S6.**
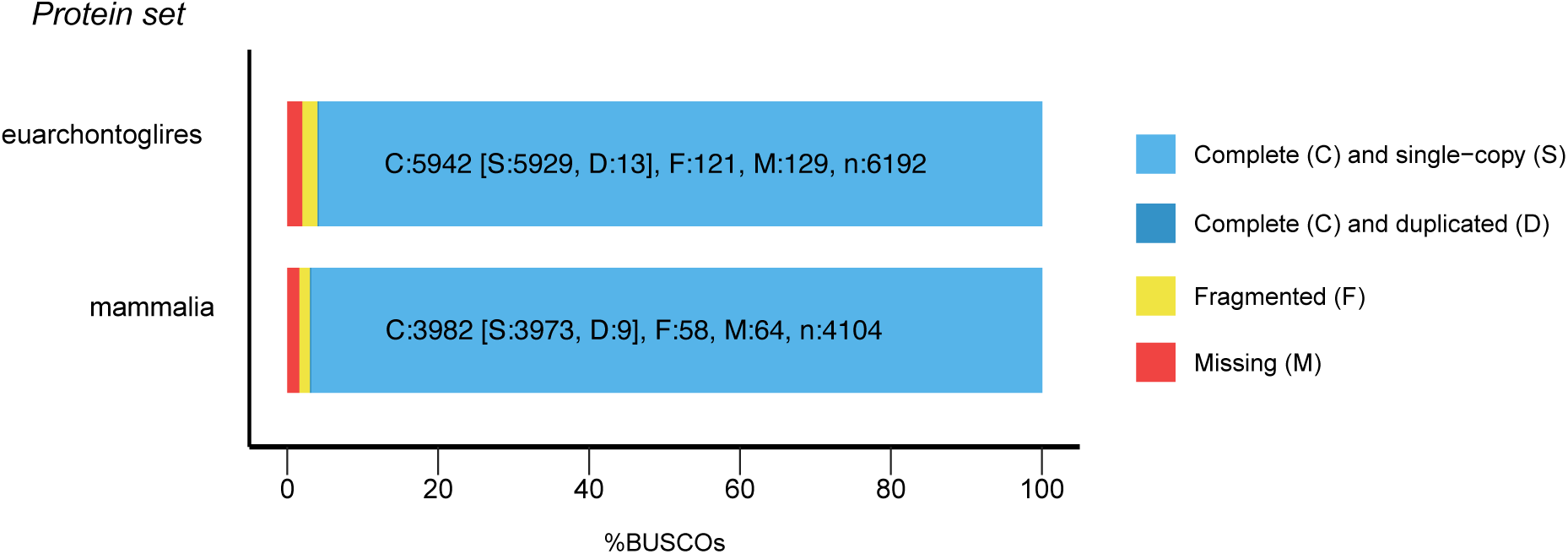
Summary assessments of annotation completeness. The bar charts for the proportion of single-copy orthologs present in the Pman2.1 version of the deer mouse genome inferred from a search against two databases (BUSCO euarchontoglires_odb9 and mammalia_odb9 databases). Light blue shows the percent of complete and single copy genes, dark blue shows percent complete and duplicated genes, yellow shows the percentage of fragmented genes and finally red shows the % of missing genes in the annotation.

**Fig. S7.**
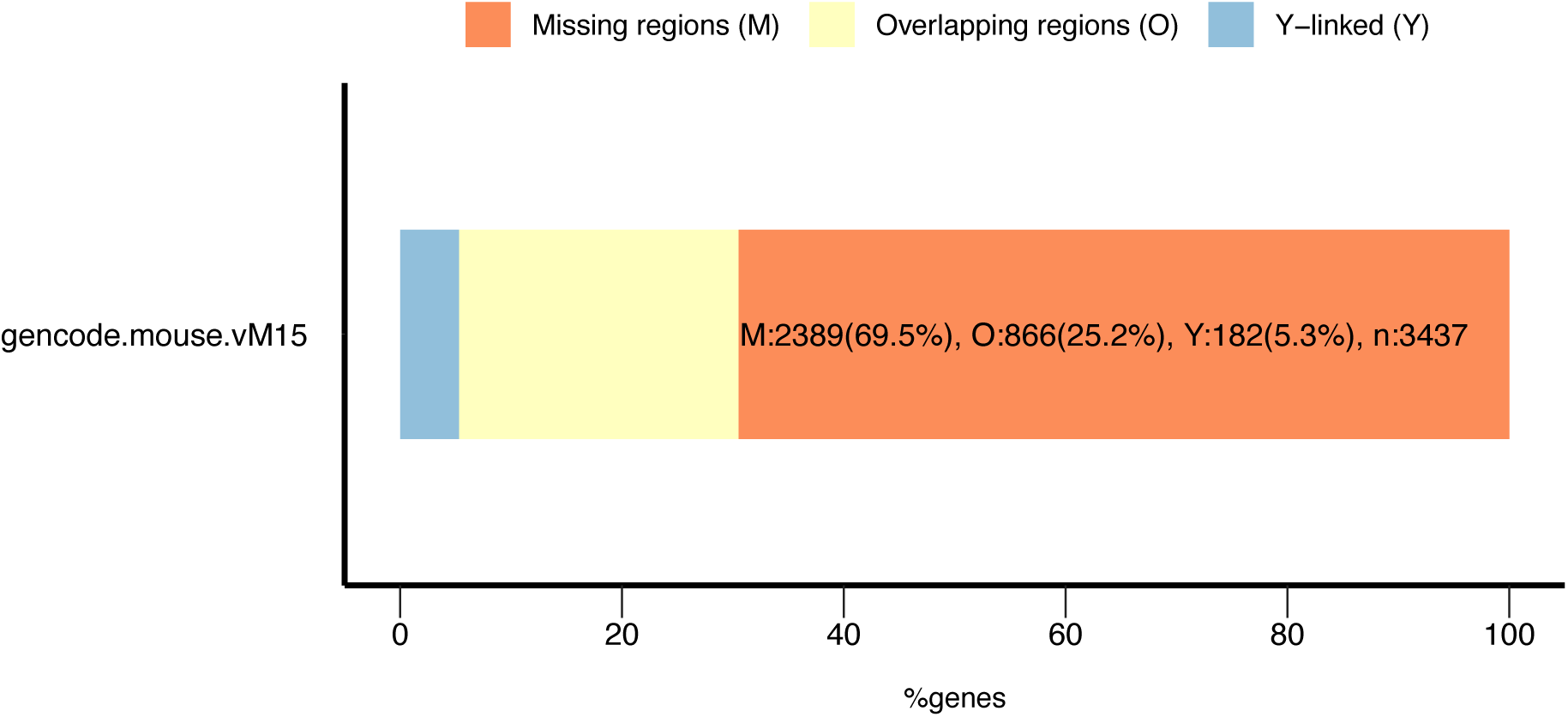
Summary assessment of mouse protein-coding genes missing in the target genome annotation. The bar charts for the proportion of protein-coding genes present in the gencode vM15 version of the laboratory mouse annotation not found in the deermouse comparative annotation. Sky blue shows the percent of genes not in the deer mouse annotation because they are on the Y chromosome, which is not covered by our assembly. It should be noted that there are 182 genes on the Y chromosome among the missing genes, while the gencode release M15 includes 183 protein-coding genes on that chromosome. This discrepancy comes from *Rbm31y*, which appears to be on the X chromosome in deer mouse. Pale yellow shows the percent of missing genes contained in overlapping genomic regions of the mouse and deer mouse genomes. Conversely, coral depicts the percent of entries in the gene list that have no overlap.

**Fig. S8.**
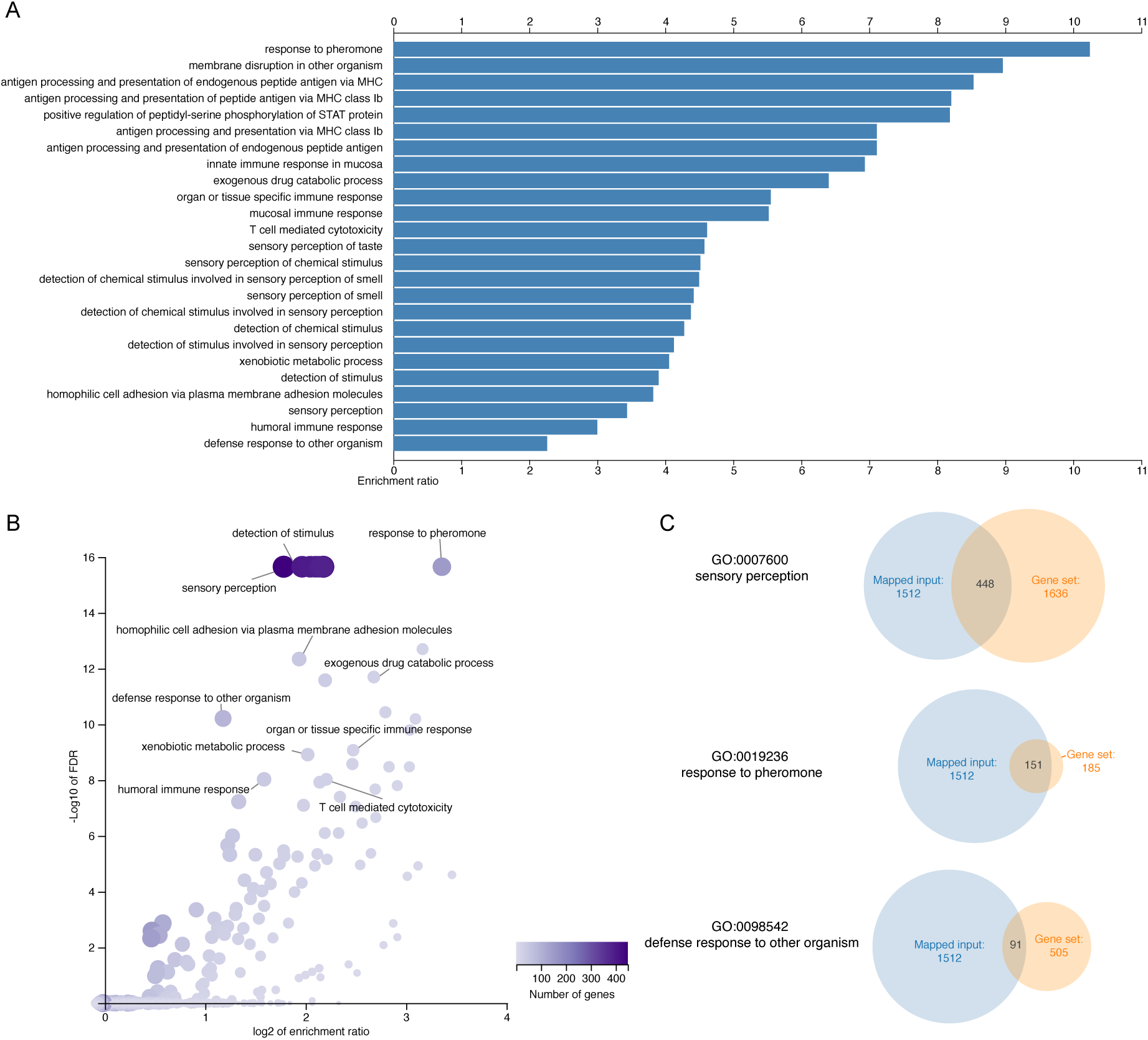
Enrichment results for genes not transferred during the comparative annotation. WebGestalt gene ontology analysis showed that genes associated with immunity/defense and chemosensory perception are enriched among the *Mus musculus* genes missing from the deer mouse annotation (**A**) The bar plot shows the biological processes associated with the genes sets that were enriched. Only the top 25 terms were annotated (*P*_FDR_ < 0.05 (Benjamini–Hochberg procedure). All bars have an FDR < 0.05. (**B**) The volcano plot shows genes sets that were enriched. Weighted set cover was applied to reduce redundancy among terms displayed. The size and color of the dots scale with the size of the gene sets. (**C**) Venn diagrams show three significantly enriched gene sets and the associated terms.

**Fig. S9.**
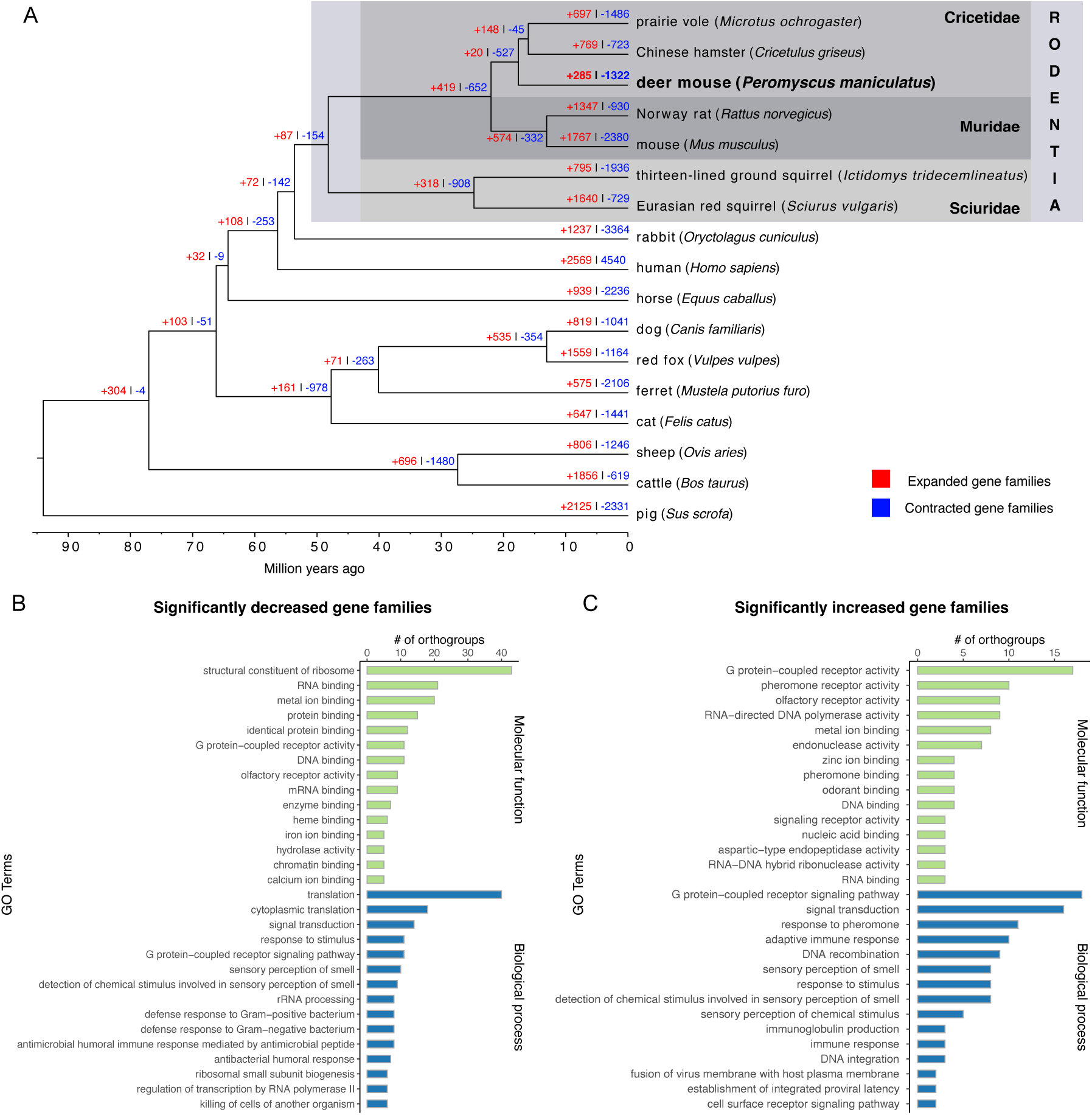
Gene expansion and contraction in the deer mouse genome. (**A**) Phylogenomic tree showing estimated divergence time and evolution of gene families for 17 mammalian species. The numbers of gene families/orthogroups that have expanded (red, +) and contracted (blue, -) after separation of the lineages are shown on the corresponding branch. Functional annotation of significantly contracted (**B**) and (**C**) expanded gene families. Only the top 15 terms are shown. Full annotation is provided as data S1. Gene ontology terms were assigned to an orthogroup based on the annotations of its members. A given GO term had to be harbored by at least two members of the orthogroups to be considered representative.

**Fig. S10.**
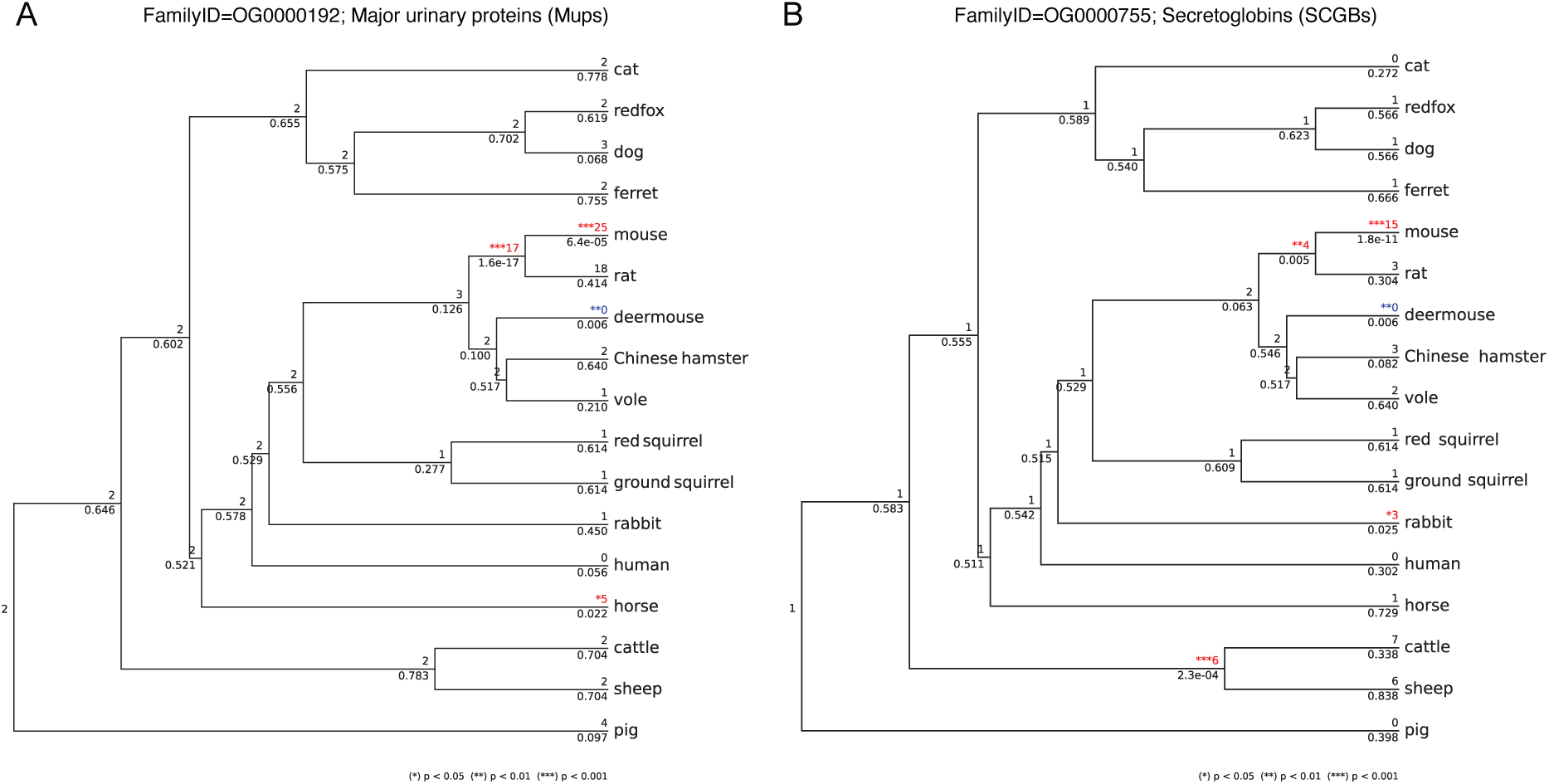
Example of gene families encoding binding proteins missing from the deer mouse genome. (**A**) Phylogenomic trees showing the evolution of genes encoding Major Urinary Proteins (MUPs) and (**B**) Secretoglobins (SCGBs). Number above branches indicate number of gene copies in the extant lineages and assignments to the ancestral nodes. Number below branches indicate p-values reported by CAFÉ. Branches with low p-values represent unusually large changes. Contractions are marked in blue and expansions are marked in red.

**Fig. S11.**
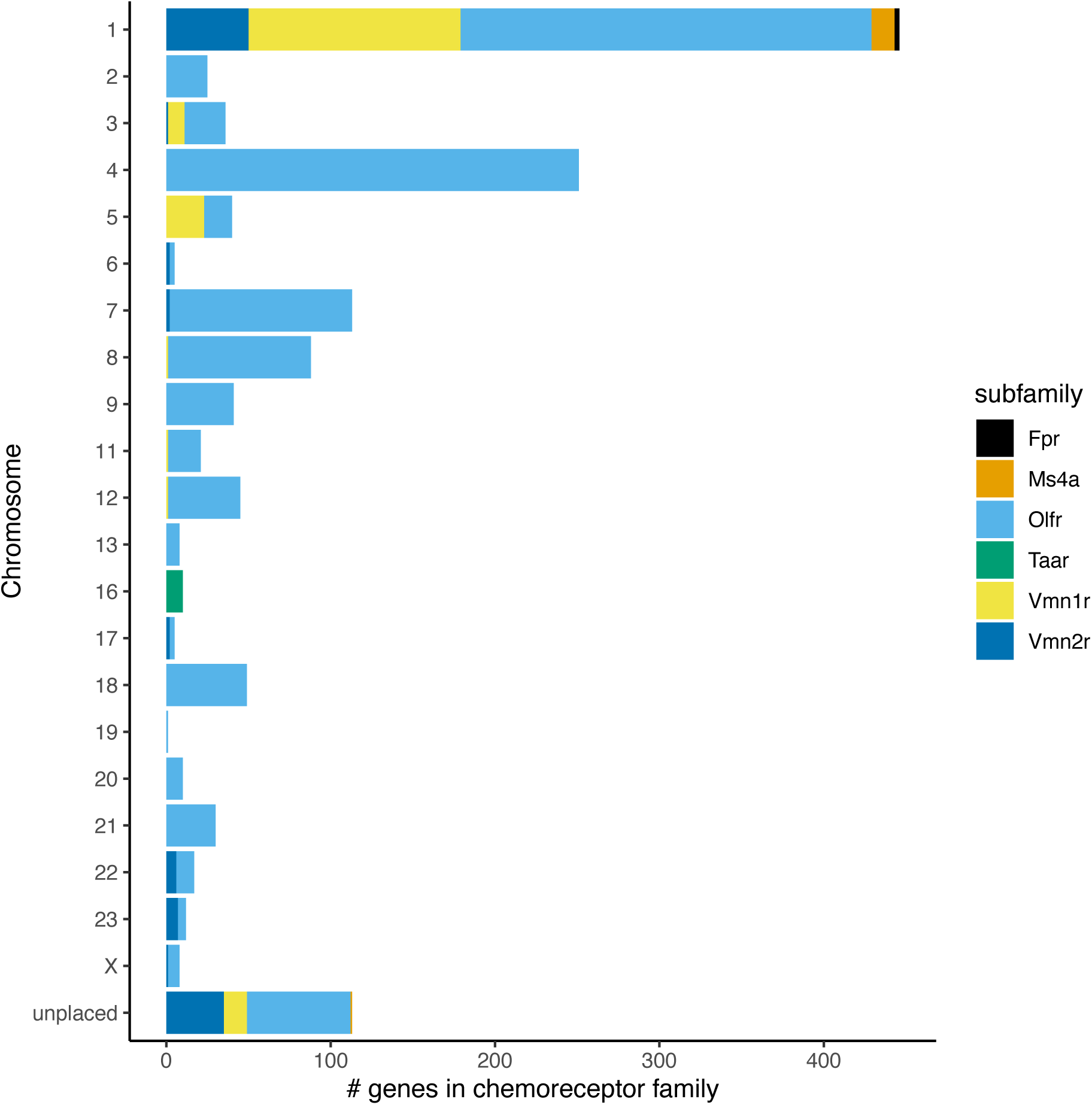
Organization of the olfactory chemosensory receptor repertoire. The bar plot shows the distribution of genes from each subfamily across the different chromosomes.

**Fig. S12.**
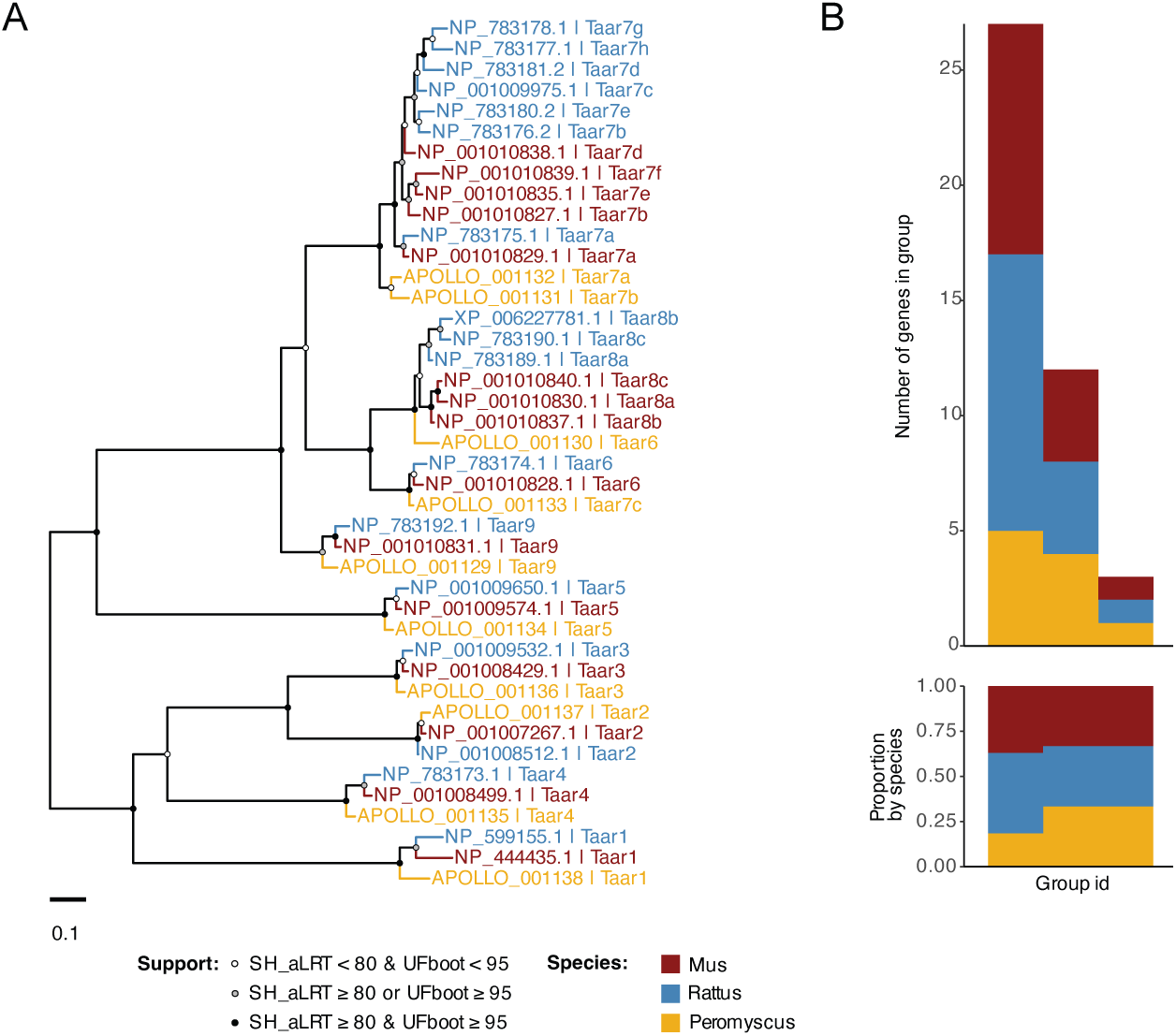
Trace amine-associated receptors repertoires. (**A**) Phylogenetic relationships between Taar genes of the laboratory mouse, rat and deer mouse. Branch support values on the maximum likelihood tree were calculated from 1000 replicates using the Shimodaira-Hasegawa-like approximate ratio test (SH_aLRT) and ultrafast bootstrapping (UFboot). Support values for branches are indicated by colored circles, with color assigned based on thresholds of branch selection for SH-aLRT (80%) and UFBoot (95%) supports, respectively. (**B**) Orthogroups intersections among the three species. The top bar plot represents the number of genes within each orthogroup intersection. The bottom bar plot represents the proportion of genes from each species within orthogroups.

**Fig. S13.**
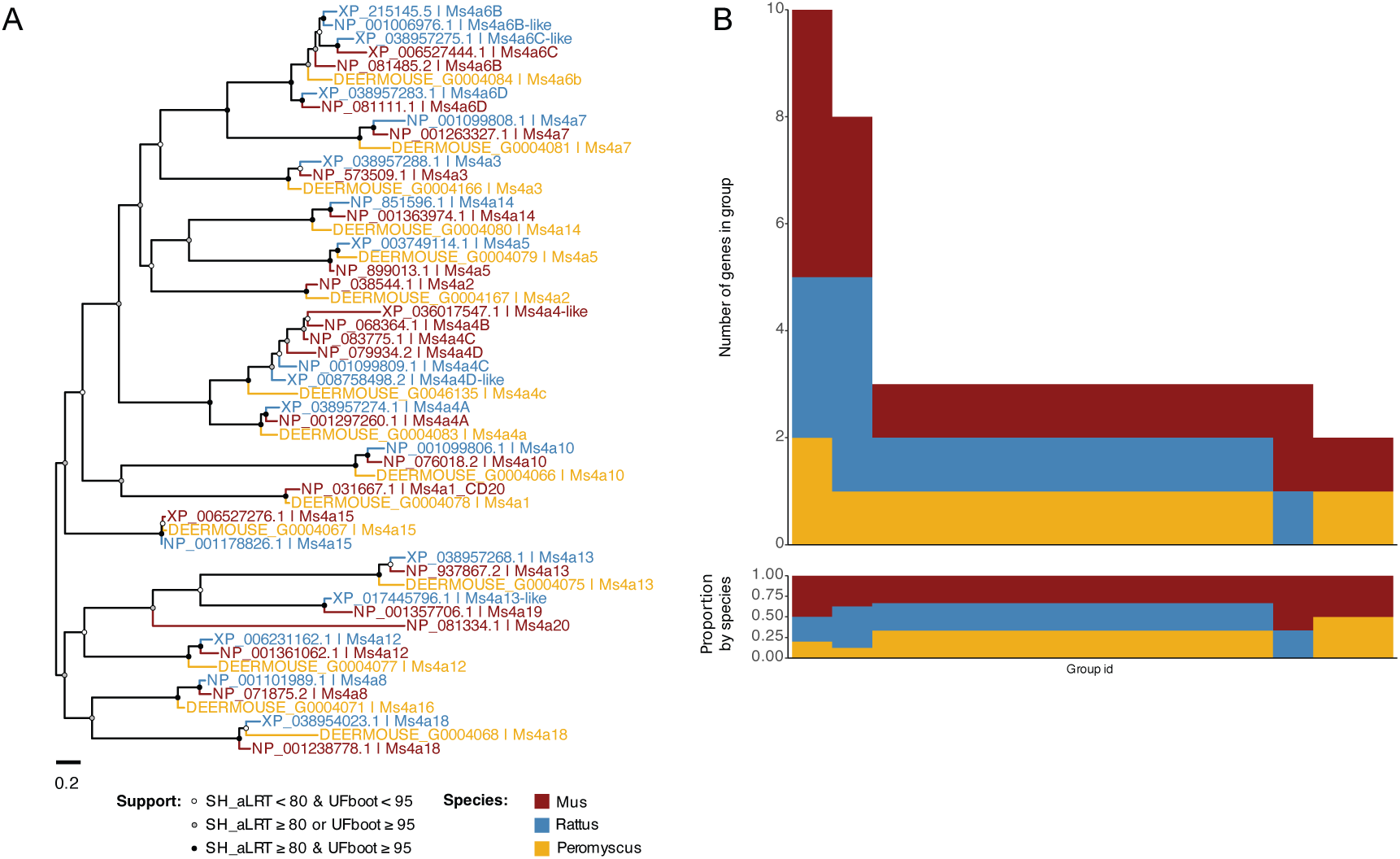
Membrane-spanning 4A (Ms4A) gene repertoires. (**A**) Phylogenetic relationships between Ms4a genes of the laboratory mouse, rat and deer mouse. Branch support values on the maximum likelihood tree were calculated from 1000 replicates using the Shimodaira-Hasegawa-like approximate ratio test (SH_aLRT) and ultrafast bootstrapping (UFboot). Support values for branches are indicated by colored circles, with color assigned based on thresholds of branch selection for SH-aLRT (80%) and UFBoot (95%) supports, respectively. (**B**) Orthogroups intersections among the three species. The top bar plot represents the number of genes within each orthogroup intersection. The bottom bar plot represents the proportion of genes from each species within orthogroups.

**Fig. S14.**
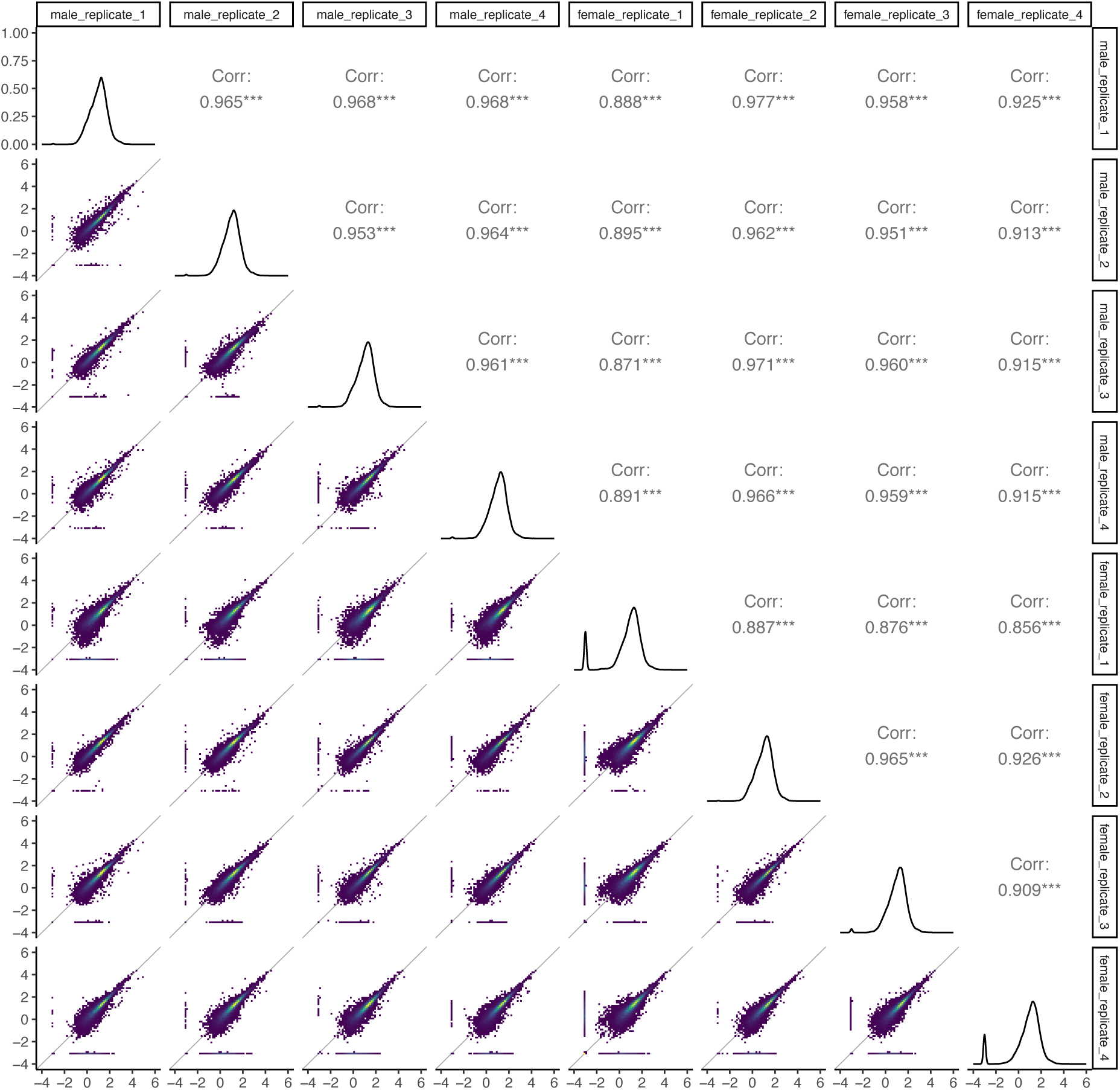
Correlations between whole olfactory mucosa (WOM) biological replicates. The lower diagonal presents pairwise scatter plots of expression estimates (log_10_ (TPM+0.001)) from WOM RNA-Seq of male and female adult deer mouse replicates. The upper diagonal shows the Spearman correlation coefficient (ρ) for each pairwise comparison.

**Fig. S15.**
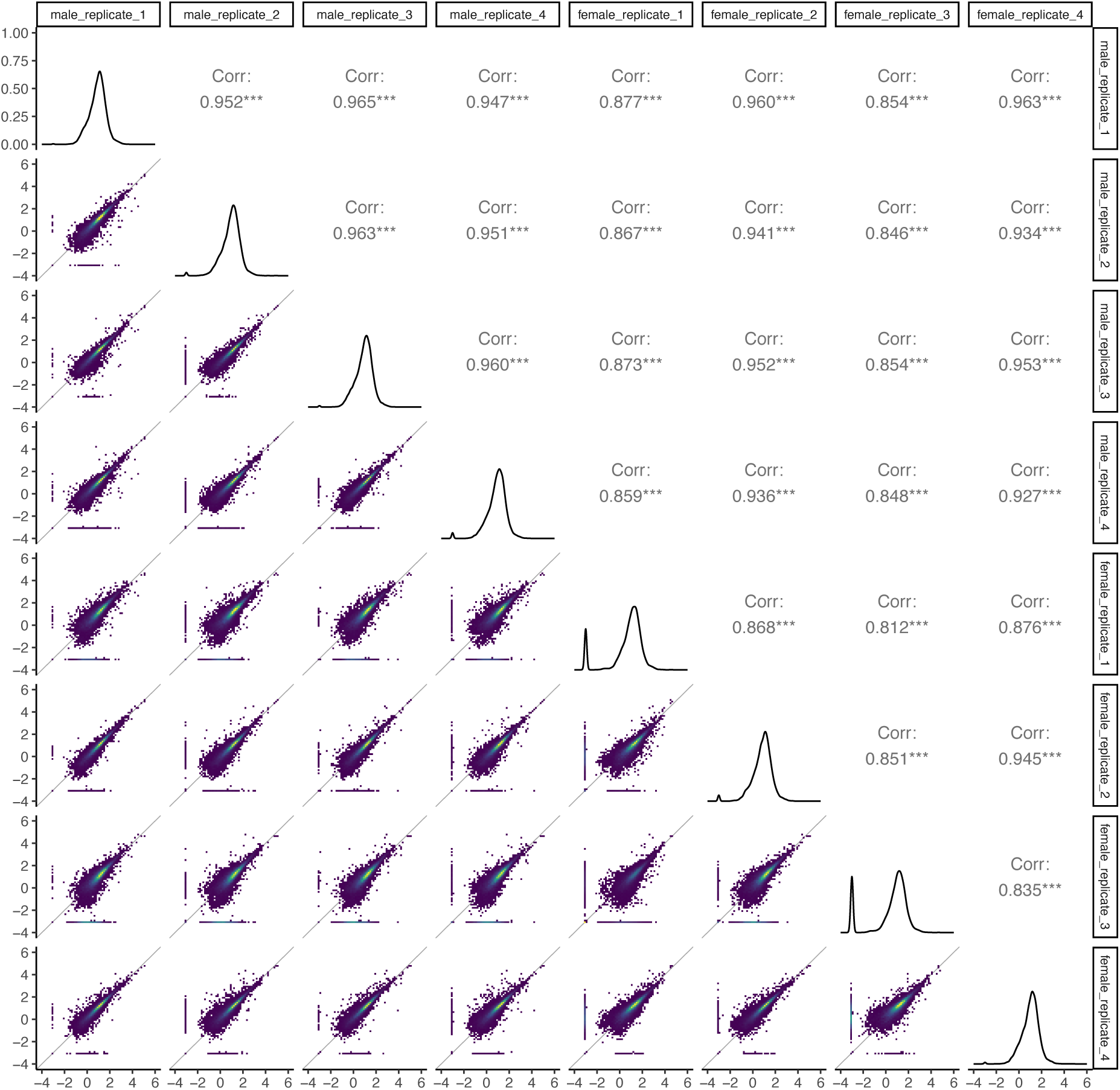
Correlations between vomeronasal organ (VNO) biological replicates. The lower diagonal presents pairwise scatter plots of expression estimates (log_10_ (TPM+0.001)) from VNO RNA-Seq of male and female adult deer mouse replicates. The upper diagonal shows the Spearman correlation coefficient (ρ) for each pairwise comparison.

**Fig. S16.**
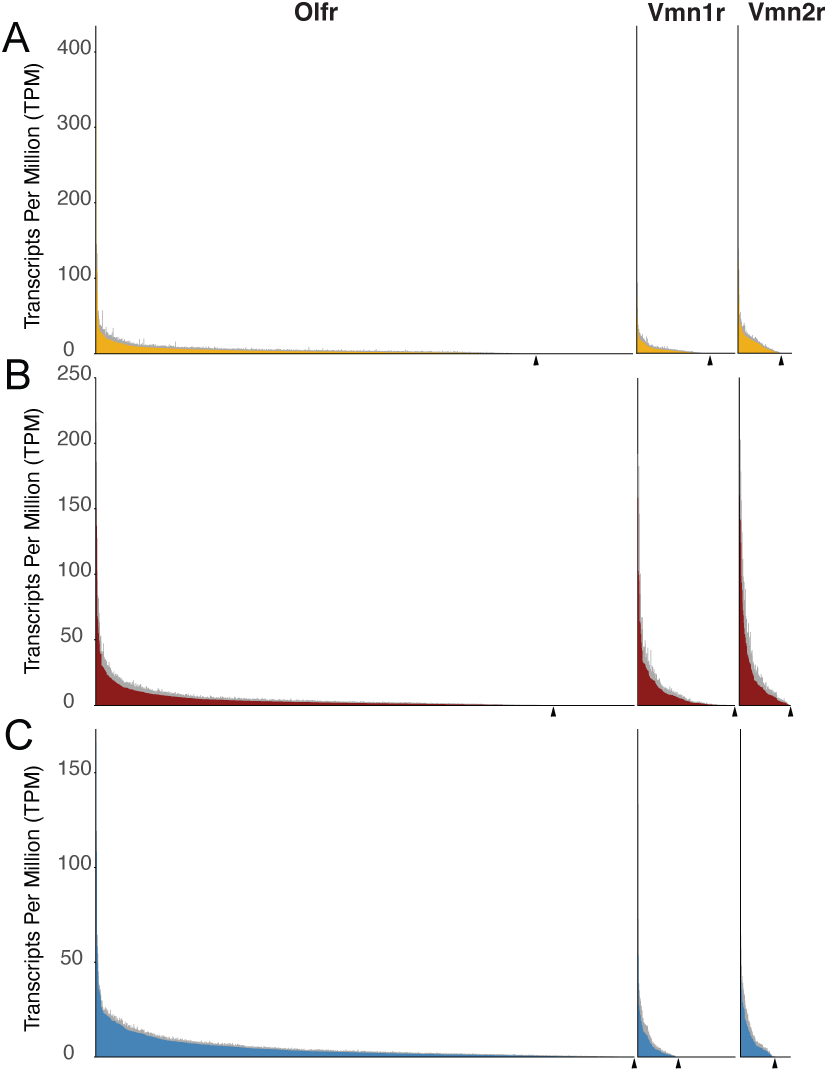
Expression profiles of nasal chemosensory receptors in rodents. Distribution of mean expression values for each receptor gene in the WOM or VNO of deer mouse (A), mouse (B), and rat (C). Expression estimates of Olfrs and Vmnrs are derived from WOM and VNO RNA-Seq data, respectively. Genes are sorted in descending order of their mean expression value (Transcript Per Million; TPM). Error bars (in grey) represent the standard deviation of the estimate from 6-8 samples (male and female samples pooled). Under the x-axis, an arrowhead indicates the position of the least abundant gene in a given family.

**Fig. S17.**
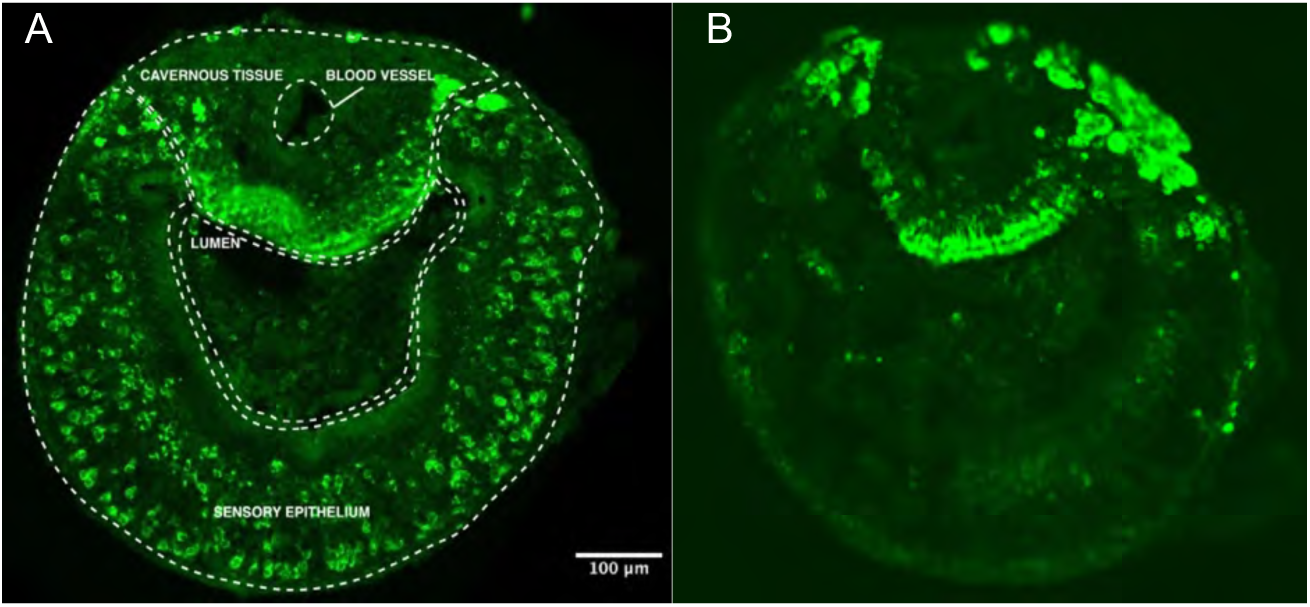
pS6 is a robust marker of activated olfactory sensory neurons. (**A**) Coronal section of the vomeronasal organ from a deer mouse harvested post-exposure to snake feces for 1 hour. Activated olfactory sensory neurons in the sensory epithelium are identified using an antibody against pS6 (240/244) and stained with a fluorescently-labeled secondary antibody (pseudo-colored in green). Green puncta in the sensory epithelium correspond to neurons responding to cues present in the stimulus. (**B**) Tissue section harvested from a deer mouse exposed to clean bedding for 1 hour as control. Low background level of pS6 activity in the sensory epithelium underlines the suitability of pS6 (240/244) as an activity-dependent marker.

**Fig. S18.**
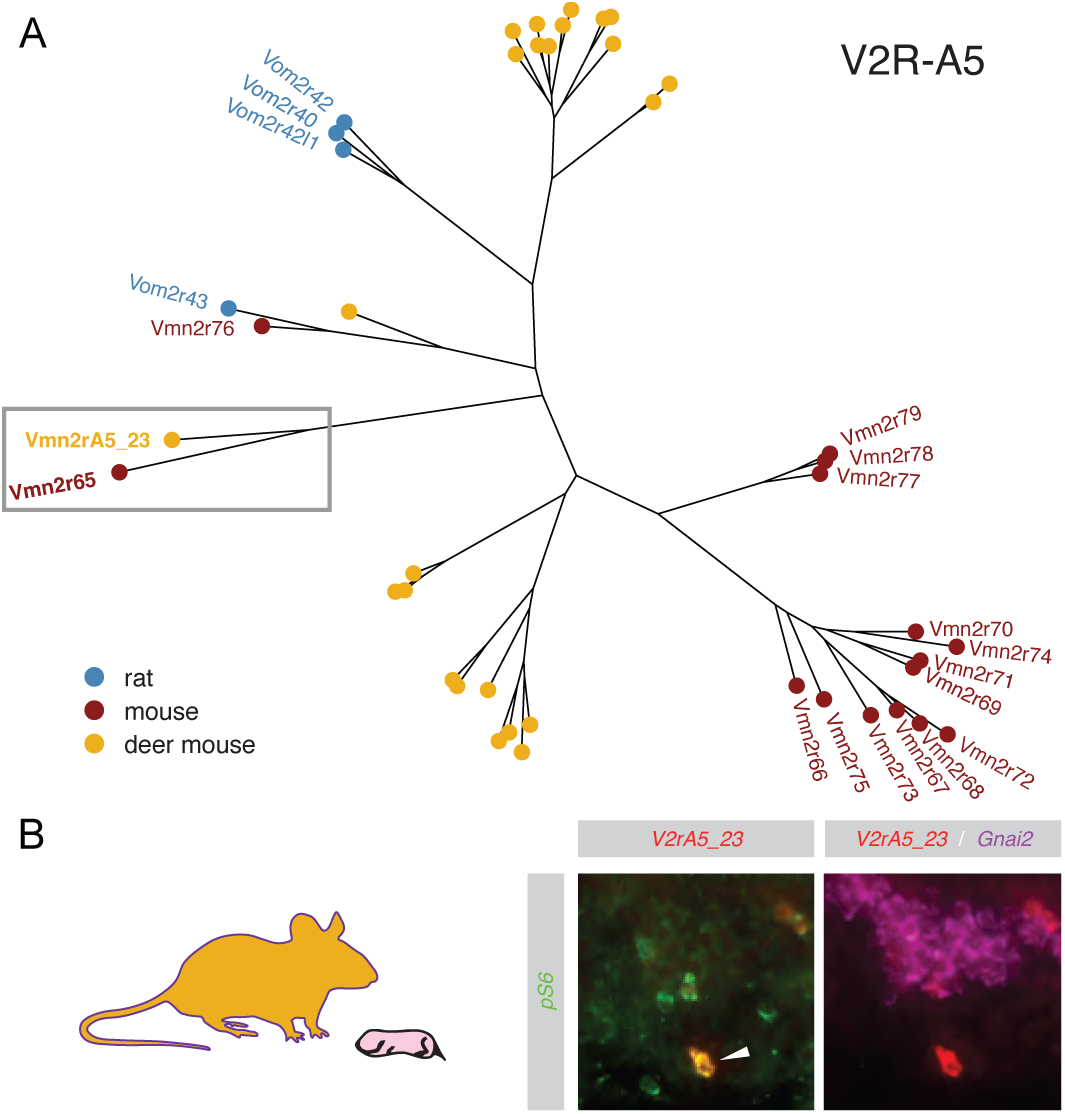
Pup cues activate V2rA5_23 in the adult VNO. (**A**) Phylogeny of clade V2R-A5 receptors. (**B**) Immunofluorescence with anti-pS6 antibody (240/244) and *in situ* hybridization with RNA probes to deer mouse V2RA5_23, the ortholog of Vmn2r65 in mouse, and Gnai2, a marker of V1R-expressing VSNs, demonstrate co-localization (arrow) in VNO neurons activated by pup cues in adult deer mouse.

**Table S1.** Overview of the sequencing data used to assemble the deer mouse genome (version HU_Pman_2.1.3).

| Library type | No. of libraries | Mean insert size, bp | Read length, bp | SRA accession | Sequencing Platform |
| --- | --- | --- | --- | --- | --- |
| Fragment | 3 | 199 | 2x100 | SRR1012303, SRR1012309, SRR1012310 | Baylor College of Medicine |
| Short jump | 3 | 500 | 2x100 | SRR1012307, SRR1012308, SRR1012311 | Baylor College of Medicine |
| Long jump | 3 | 3000 | 2x100 | SRR1012304, SRR1012305, SRR1012306 | Baylor College of Medicine |
| Fosmid jump | 1 | 40000 | 2x100 | SRR10402715 | Broad Institute; *This study* |

**Table S2.**
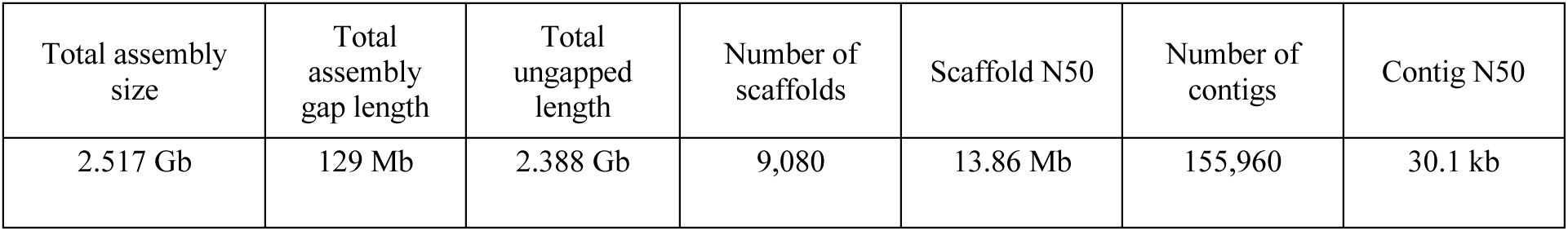
Statistics for the ALLPATH-LG assembly.

| Total assembly size | Total assembly gap length | Total ungapped length | Number of scaffolds | Scaffold N50 | Number of contigs | Contig N50 |
| --- | --- | --- | --- | --- | --- | --- |
| 2.517 Gb | 129 Mb | 2.388 Gb | 9,080 | 13.86 Mb | 155,960 | 30.1 kb |

**Table S3.** Summary of statistics for markers used to generate the consensus assembly of HU_Pman2.1.3 pseudo-chromosomes.

|  | Anchored | Oriented | Unplaced |
| --- | --- | --- | --- |
| Number of unique markers | 62,992 | 62,830 | 396 |
| Markers per Mb | 25.9 | 25.9 | 5.2 |
| Scaffold N50 | 59 | 59 | 0 |
| Scaffolds | 585 | 447 | 8,505 |
| Scaffolds with 1 marker | 126 | 0 | 173 |
| Scaffolds with 2 markers | 67 | 61 | 33 |
| Scaffolds with 3 markers | 19 | 16 | 12 |
| Scaffolds with 4 or more markers | 373 | 370 | 14 |
| Total number of bases | 2,436,387,942 | 2,427,689,157 | 75,987, 719 |
| Percentage of sequenced bases | 97.0% | 96.6% | 3.0% |

**Table S4.** Comparative annotation statistics. This table reports the number of genes in the original annotation set (mm10 gencode v15) found or missing in the target genome.

| Gene Class | Complete | Missing | Completeness |
| --- | --- | --- | --- |
| 3prime_overlapping_ncRNA | 2 | 0 | 100.0% |
| antisense_RNA | 2516 | 130 | 95.1% |
| bidirectional_promoter_lncRNA | 120 | 4 | 96.8% |
| IG_C_gene | 8 | 5 | 61.5% |
| IG_C_pseudogene | 1 | 0 | 100.0% |
| IG_J_gene | 10 | 2 | 83.3% |
| IG_LV_gene | 1 | 3 | 25.0% |
| IG_V_gene | 17 | 201 | 7.8% |
| IG_V_pseudogene | 11 | 143 | 7.1% |
| lincRNA | 4060 | 1022 | 79.9% |
| macro_lncRNA | 1 | 0 | 100.0% |
| miRNA | 1248 | 959 | 56.5% |
| misc_RNA | 419 | 144 | 74.4% |
| polymorphic_pseudogene | 42 | 35 | 54.5% |
| processed_pseudogene | 2405 | 6211 | 27.9% |
| processed_transcript | 689 | 80 | 89.6% |
| protein_coding | 18529 | 3472 | 84.2% |
| pseudogene | 45 | 29 | 60.8% |
| ribozyme | 9 | 13 | 40.9% |
| rRNA | 177 | 177 | 50.0% |
| scaRNA | 30 | 21 | 58.8% |
| scRNA | 1 | 0 | 100.0% |
| sense_intronic | 285 | 22 | 92.8% |
| sense_overlapping | 26 | 1 | 96.3% |
| snoRNA | 386 | 1121 | 25.6% |
| snRNA | 527 | 858 | 38.1% |
| sRNA | 2 | 0 | 100.0% |
| TEC | 2719 | 297 | 90.2% |
| TR_C_gene | 5 | 3 | 62.5% |
| TR_J_gene | 62 | 2 | 96.9% |
| TR_J_pseudogene | 9 | 1 | 90.0% |
| TR_V_gene | 61 | 83 | 42.4% |
| TR_V_pseudogene | 20 | 13 | 60.6% |
| transcribed_processed_pseudogene | 136 | 110 | 55.3% |
| transcribed_unitary_pseudogene | 12 | 0 | 100.0% |
| transcribed_unprocessed_pseudogene | 101 | 125 | 44.7% |
| unitary_pseudogene | 18 | 0 | 100.0% |
| unprocessed_pseudogene | 527 | 2009 | 20.8% |
| **TOTAL** | **35237** | **17296** | **67.1%** |

**Data S1.**

Complete set of CAFE_gene_families_with functions.

**Data S2.**

Complete set of orthogroups with conserved receptors with annotations.

